# O-GlcNAc Signaling Increases Neuron Regeneration Through One-Carbon Metabolism in *Caenorhabditis elegans*

**DOI:** 10.1101/2023.03.05.531166

**Authors:** Dilip Kumar Yadav, Andrew C Chang, Christopher V Gabel

**Affiliations:** Physiology and Biophysics; Pharmacology and Experimental Therapeutics; Neurophotonics Center, Boston University Chobanian & Avedisian School of Medicine, Boston MA 02118 USA

## Abstract

Cellular metabolism plays an essential role in the regrowth and regeneration of a neuron following physical injury. Yet, our knowledge of the specific metabolic pathways that are beneficial to neuron regeneration remains sparse. Previously, we have shown that modulation of O-linked β-N-acetylglucosamine (O-GlcNAc), a ubiquitous post-translational modification that acts as a cellular nutrient sensor, can significantly enhance *in vivo* neuron regeneration. Here we define the specific metabolic pathway by which mutation of the O-GlcNAc transferase (*ogt-1)* increases regenerative outgrowth. Performing *in vivo* laser axotomy and measuring subsequent regeneration of individual neurons in *C. elegans*, we find that the *ogt-1* mutation increases regeneration by diverting the metabolic flux of enhanced glycolysis towards one carbon metabolism (OCM) and the downstream transsulfuration metabolic pathway (TSP). These effects are abrogated by genetic and/or pharmacological disruption of OCM or the serine synthesis pathway (SSP) that links OCM to glycolysis. Testing downstream branches of this pathway, we find that enhanced regeneration is dependent only on the vitamin B12 independent shunt pathway. These results are further supported by RNA-sequencing that reveals dramatic transcriptional changes, by the *ogt-1* mutation, in the genes involved in glycolysis, OCM, TSP and ATP metabolism. Strikingly, the beneficial effects of the *ogt-1* mutation can be recapitulated by simple metabolic supplementation of the OCM metabolite methionine in wild-type animals. Taken together, these data unearth the metabolic pathways involved in the increased regenerative capacity of a damaged neuron in *ogt-1* animals and highlight the therapeutic possibilities of OCM and its related pathways in the treatment of neuronal injury.

**Abstarct Figure.**
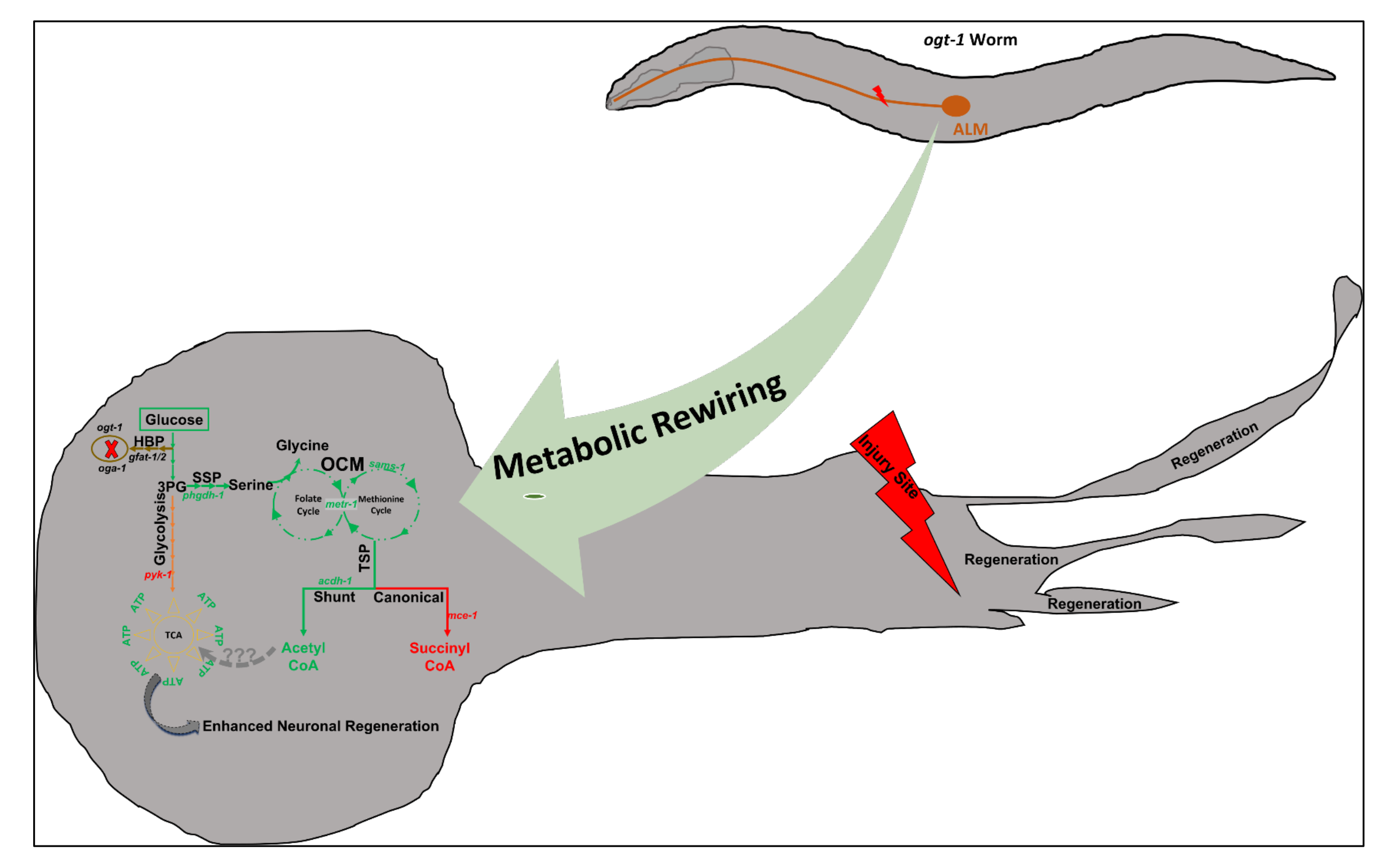
Metabolic pathways involved in the enhanced neuronal regeneration in *ogt-1* animals: The green highlighted pathway illustrates the metabolic rewiring in *ogt-1* mutant animals supporting enhanced axonal regeneration of injured neurons *in vivo*.

## INTRODUCTION

To regenerate efficiently, a damaged neuron must undergo molecular and metabolic rearrangement to induce and endure a range of complex cellular processes (Mahar and Cavalli 2018, Taub, Awal et al. 2018, Yang, Wang et al. 2020). These processes are extremely metabolically challenging, energy demanding and critical for the regenerative capacity of a neuron (Byrne, Walradt et al. 2014, Cartoni, Norsworthy et al. 2016, He and Jin 2016, Yang, Wang et al. 2020). The importance of metabolic pathways, particularly in neuronal regeneration including the insulin-signaling pathway, energy metabolism, and mitochondrial function have been reported in research articles by several groups (Byrne, Walradt et al. 2014, Cartoni, Norsworthy et al. 2016, Han, Baig et al. 2016, Han, Xie et al. 2020). Nonetheless critical questions remain as to the alterations in cellular metabolism and energy production in a damaged and regenerating neuron and how these processes might be exploited for therapeutic benefits.

In a previous study our group demonstrated that perturbation in O-linked β-N-acetylglucosamine (O-GlcNAc) signaling, a post-translational modification of serine and threonine that is known to act as a nutrient sensor, substantially increased axonal regeneration in *Caenorhabditis elegans* (*C. elegans*) (Taub, Awal et al. 2018). Carrying out *in vivo* laser axotomies, we demonstrated that a reduction of O-GlcNAc levels, due to mutation of the O-GlcNAc transferase (*ogt-1*), induces the AKT-1 branch of the insulin-signaling pathway to utilize glycolysis and significantly enhanced neuronal regeneration. Inhibition of the glycolytic pathway through RNAi knockdown of phosphoglycerate kinase (*pgk-1*) or loss of function of phosphofructokinase-1.1 (*pfk-1.1*) specifically suppressed *ogt-1* enhanced regeneration but did not alter wild-type regeneration (Taub, Awal et al. 2018). Furthermore, supplementation with glucose in wild type animals is sufficient to increase axonal regeneration after axotomy (Taub, Awal et al. 2018). These observations established the importance of increased glycolytic metabolism to control and enhance neuronal regeneration.

To date key questions, remain as to what specific metabolic pathways are stimulated in the *ogt-1* mutant background and what cellular processes are augmented to increase regenerative capacity. Numerous reports suggest that increased glycolysis averts metabolic flux towards one carbon metabolism (OCM) to regulate numerous biological processes including molecular reprogramming, immunological functions as well as neuronal development and function (Iskandar, Rizk et al. 2010, Konno, Asai et al. 2017, Yu, Wang et al. 2019). In addition, studies have reported the importance of metabolic amendments of OCM, the serine synthesis pathway (SSP) and the transsulfuration pathway (TSP) in neuronal development, structure, function, and regeneration (Iskandar, Rizk et al. 2010, Bonvento and Bolaños 2021, Lam, Kervin et al. 2021, Chen, Calandrelli et al. 2022). Measuring neuronal regeneration in *C. elegans* following laser axotomy under genetic, pharmacological, and metabolic perturbations, we demonstrate that *ogt-1* mutation in fact diverts glycolytic flux to OCM *via* the SSP and that functional OCM and SSP are both essential for enhanced neuronal regeneration in *ogt-1* animals. From there, we observed that metabolic flux from OCM through the TSP resulting in cystathionine metabolism into Acetyl-CoA *via* the vitamin B12 independent shunt pathway is also critical to *ogt-1* regeneration. Taken together our results illustrate how *ogt-1* acts as a major regulator of metabolic flux to orchestrate and maximize the regenerative response in a damaged neuron and suggest that OCM and its related pathways could serve as a potent neurotherapeutic target.

## RESULTS

### 1. Enhanced glycolysis pathway is sufficient for increased neuronal regeneration in *ogt-1* animals

Following our previous study, we sought to verify that increased glycolytic flux is the main mechanism of increased neuronal regeneration in the *ogt-1* mutant background. Performing laser axotomy on individual neurons and measuring regenerative outgrowth after 24 h, we found that *ogt-1* mutation increases the neuronal regeneration in ALM and PLN neurons in *C. elegans* (fig 1A-1B, Table-S1. and supp fig 1A, Table-S1). Reduced O-GlcNAc levels due to the *ogt-1* mutation will effectively block the metabolic flux into the Hexosamine Biosynthesis Pathway (HBP) diverting metabolites towards glycolysis (Yi, Clark et al. 2012, Jóźwiak, Forma et al. 2014, Kim, Nakayama et al. 2018). We recapitulated this effect by knocking down Glutamine-Fructose 6-phosphate Amino Transferase (*gfat-1* and *gfat-2*) using neuron specific RNAi. *gfat-1* and *gfat-2*, orthologs of the human, glutamine--fructose-6-phosphate transaminase 1 (*GFPT1*), catalyze the very first and rate limiting step of HBP (fig. 1A). We found that knocking down either *gfat-1* or *gfat-2* significantly increases the regeneration of ALM neurons in *C. elegans* compared to RNAi control in wild type (fig.1C, Table-S1). Earlier we have reported that genetic inhibition of the glycolytic enzymes phosphoglycerate kinase (*pgk-1*) and phosphofructokinase-1.1 (*pfk-1*), both of which work in early steps of the glycolysis pathway, suppresses *ogt-1* neuronal regeneration (Supp fig 1B; and Taub et al). Reports suggest O-GlcNAc levels regulate the expression and activity of pyruvate kinase, PKM1/2, encoded by *pyk-1* in *C. elegans*, which catalyzes the final step of glycolysis to produce pyruvate from phosphoenolpyruvate (Wang, Liu et al. 2017, Bacigalupa, Bhadiadra et al. 2018, Yu, Teoh et al. 2019). *C. elegans* has two orthologs of mammalian PK, *pyk-1* and *pyk-2,* with *pyk-1* expression primarily in neurons including the ALM and PLM neurons and *pyk-2* showing limited neuron expression (Hammarlund, Hobert et al. 2018) (supp fig. 1A and 1B). We found that knock down of the *C. elegans* ortholog, *pyk-1, via* neuron specific RNAi, does not affect regeneration in the *ogt-1* mutant but significantly increases regeneration in WT (fig. 1A, 1D, Table-S1), effectively phenocopying *ogt-1*. Furthermore, by performing *pyk-1* activity assay in whole worm lysate we observed that over all *pyk-1* activity is significantly down in *ogt-1* worms (fig. 1E).

**Fig.1. Enhanced.**
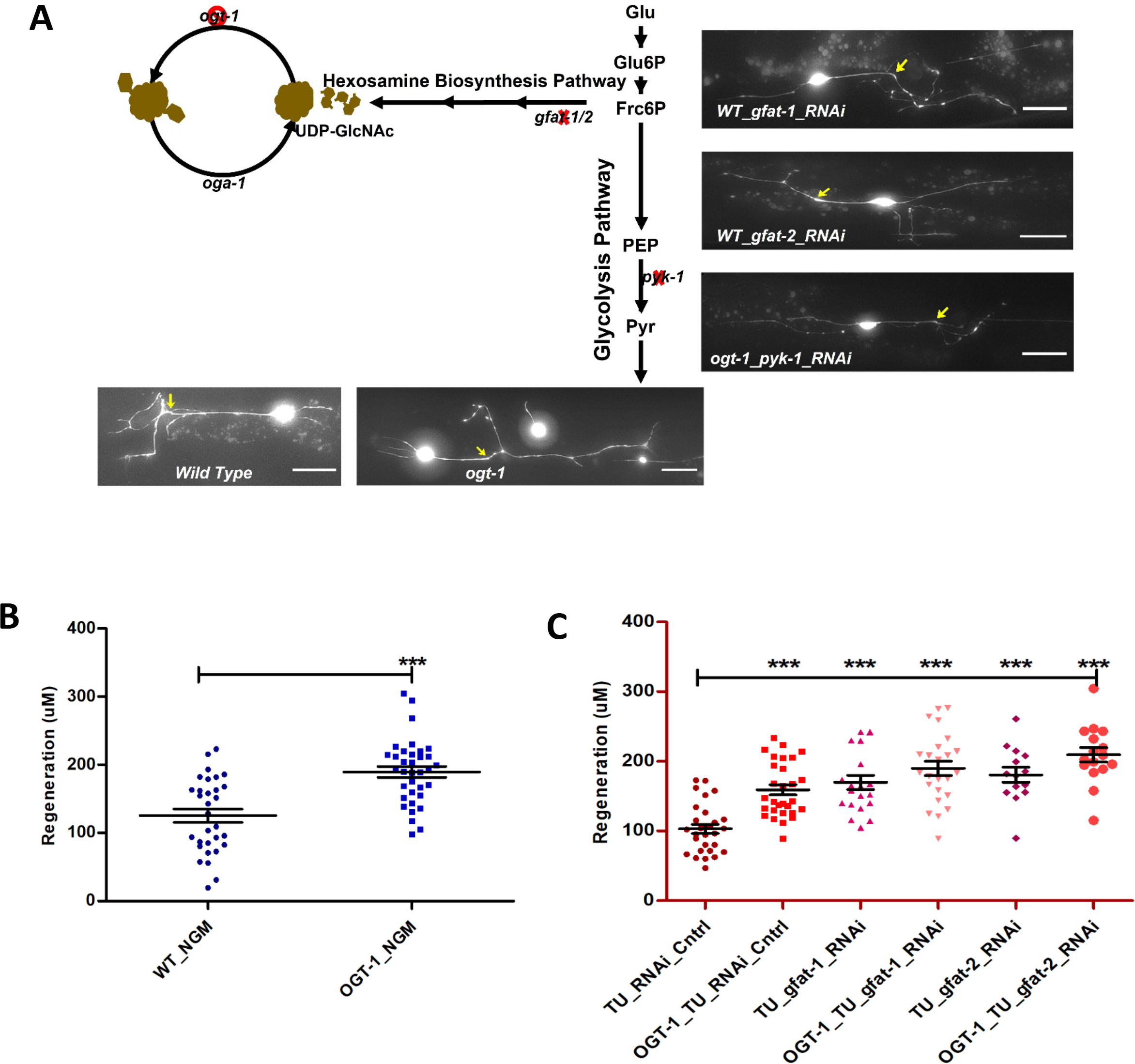
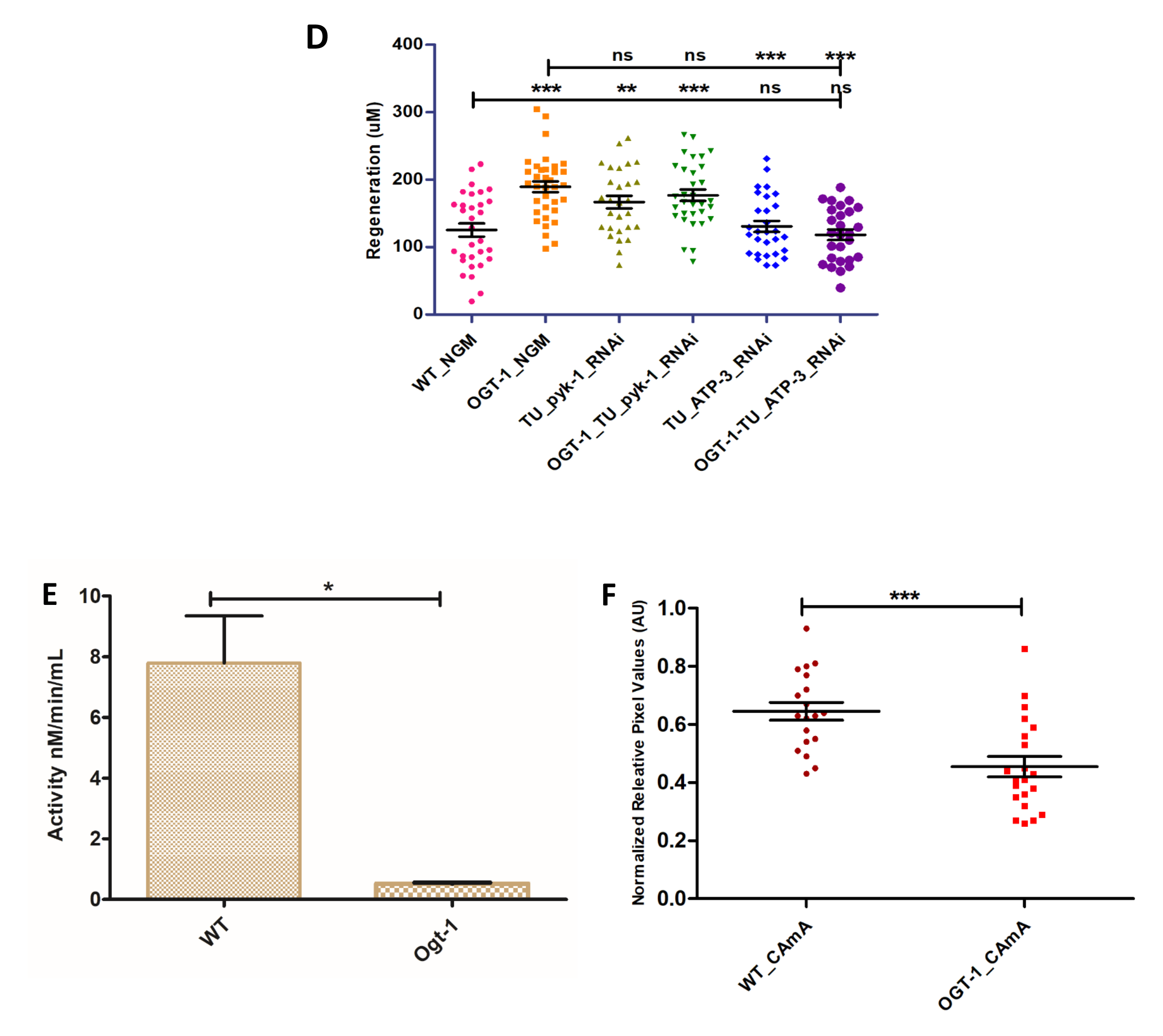
glycolysis is sufficient to support neuronal regeneration. **(A)** Schematic diagram showing the hexosamine synthesis pathway linking glycolysis and *ogt-1* function, and the effect of *ogt-1* mutation and *gfat-1/gfat-2* and *pyk-1* RNAi knockdown on regenerating neurons imaged at 24 h (arrow indicates the point of injury). **(B)** 24 h regeneration data of WT and *ogt-1* mutant worms. **(C)** 24 h regeneration data of control and *gfat-1/gfat-2* RNAi experiments. **(D)** 24 h regeneration data of control and RNAi experiment for *pyk-1* and *atp-3*. **(E)** pyk-1 activity measured in WT and *ogt-1* animal whole lysate using Pyruvate Kinase (PK) Assay Kit (Abcam, cat# Ab83432). **(F)** Relative amount of ATP measured using a FRET-based ATP sensor. AU; Arbitrary Unit, scale bar = ∼10uM, all data shown in ±SEM, analytical methods student t-test and One Way ANOVA, *pValue <0.05, **pValue <0.01, ***pValue <0.001.

In Taub *et al* we present strong evidence that enhanced glycolytic metabolism is a key element of the increased regeneration in *ogt-1* animals. To further investigate if the energy production is critical for these effects, we performed neuron specific RNAi knockdown of *atp-3*, an ortholog of human ATP5PO (ATP synthase peripheral stalk subunit OSCP) predicted to have proton-transporting ATP synthase activity. *atp-3* knockdown abrogated the *ogt-1* mediated enhanced regeneration but has no effect on regeneration in WT (Fig. 1D, Table-S1). However, these effects did not translate to whole animal ATP level measurements. Employing a FRET-based transgenic fluorescence ATP sensor (as described earlier in Soto et al, 2020) (Fig. 1F, Table-S1) as well as ATP measurements in whole worm lysate, we found that ATP levels were significantly lower in *ogt-1* than WT worms (supp Fig. 1C). In addition, we assessed whole animal ATP utilization measuring pyrophosphate (PPi) levels as an indirect indication of ATP hydrolysis but found no measurable difference between WT and *ogt-1* worms (supp Fig. 1D). Taken as a whole, these results confirm that increased flux through the majority of the glycolytic pathway and neuron specific ATP production is indeed important for *ogt-1* mediated enhanced regeneration, but that a complex interaction of metabolic pathways beyond that of canonical glycolysis may be involved specifically within the damaged and regenerating neuron.

### 2. Gene expression analysis reveals the involvement of One Carbon Metabolism (OCM) and its offshoot pathways in enhanced neuron regeneration in *ogt-1* animals

To identify additional genes and pathways involved in the enhanced regeneration of the *ogt-1* mutant, we took an unbiased approach measuring differential gene expression *via* RNAseq analysis in WT and *ogt-1* mutants. We first executed RNAseq analysis from RNA isolated from whole animals and identified a substantial number of differentially expressed genes (DEGs) in *ogt-1* compared to WT (Fig. 2A, Table-S2). Gene ontology (GO) and Kegg pathway classification analysis of DEGs identified metabolic processes such as carbohydrate, lipid, amino acids, and nucleotide metabolism as the most enriched biological processes (Fig. 2B-2C, Table-S2). In addition, cell membrane, cargo transport, nutrient reservoir and energy metabolism are also enriched in *ogt-1* (Fig. 2B-2C, Table-S2). Kegg metabolic pathway enrichment analysis revealed the enrichment of xenobiotics, drug metabolism along with glutathione metabolism, energy metabolism, amino acid, and nitrogen metabolic pathways (Fig. 2D, Table-S2). GO molecular function analysis highlights the nutrient reservoir, glutathione and s-adenosyl methionine (SAM) dependent molecular functions (Fig. 2E, Table-S2). The enrichment of amino acid, nucleotide, glutathione, and SAM dependent metabolic pathways indicates a possible role of one carbon metabolism (OCM) and its offshoot pathways in *ogt-1* mutant mediated regeneration.

**Fig.2.**
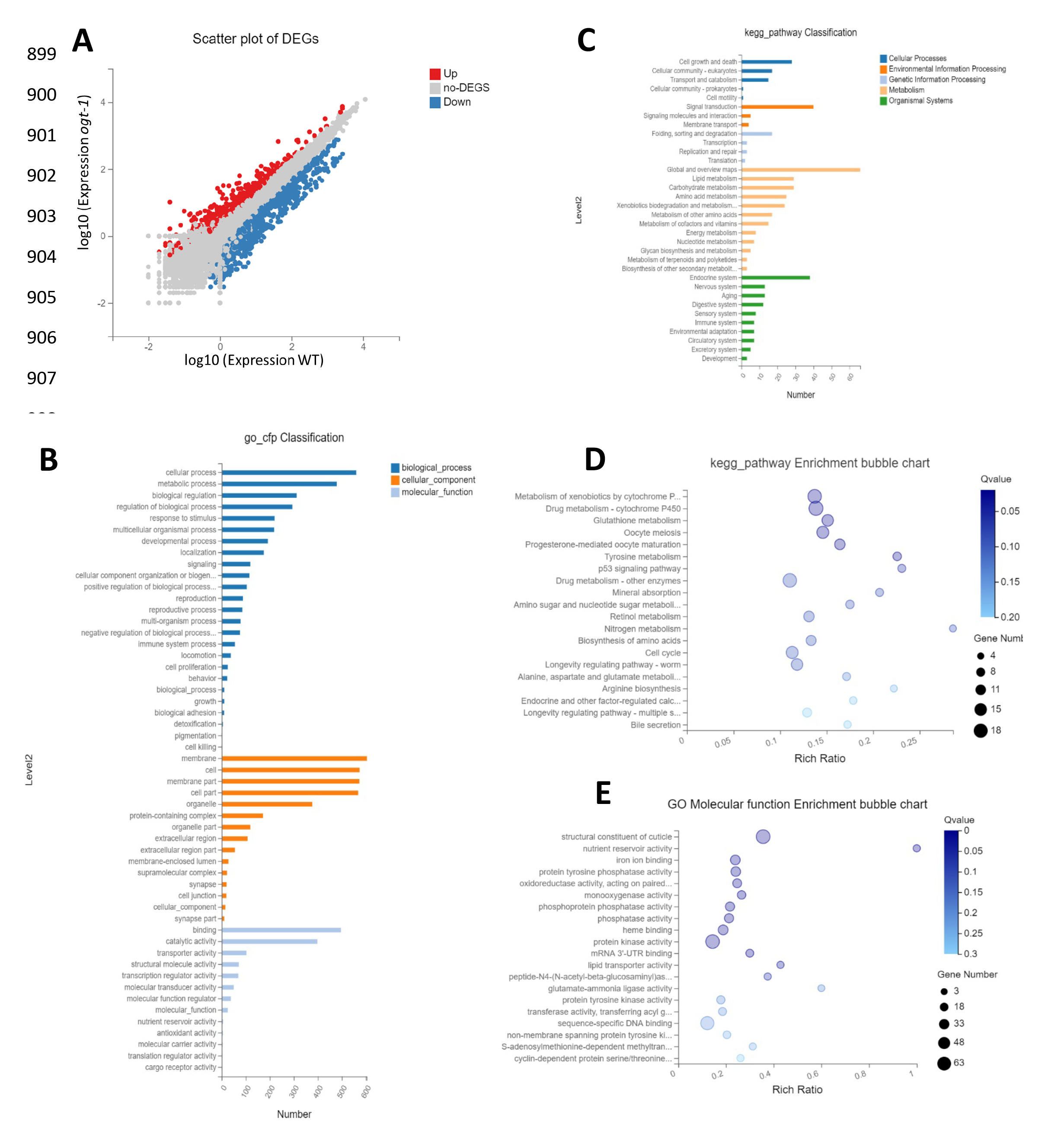
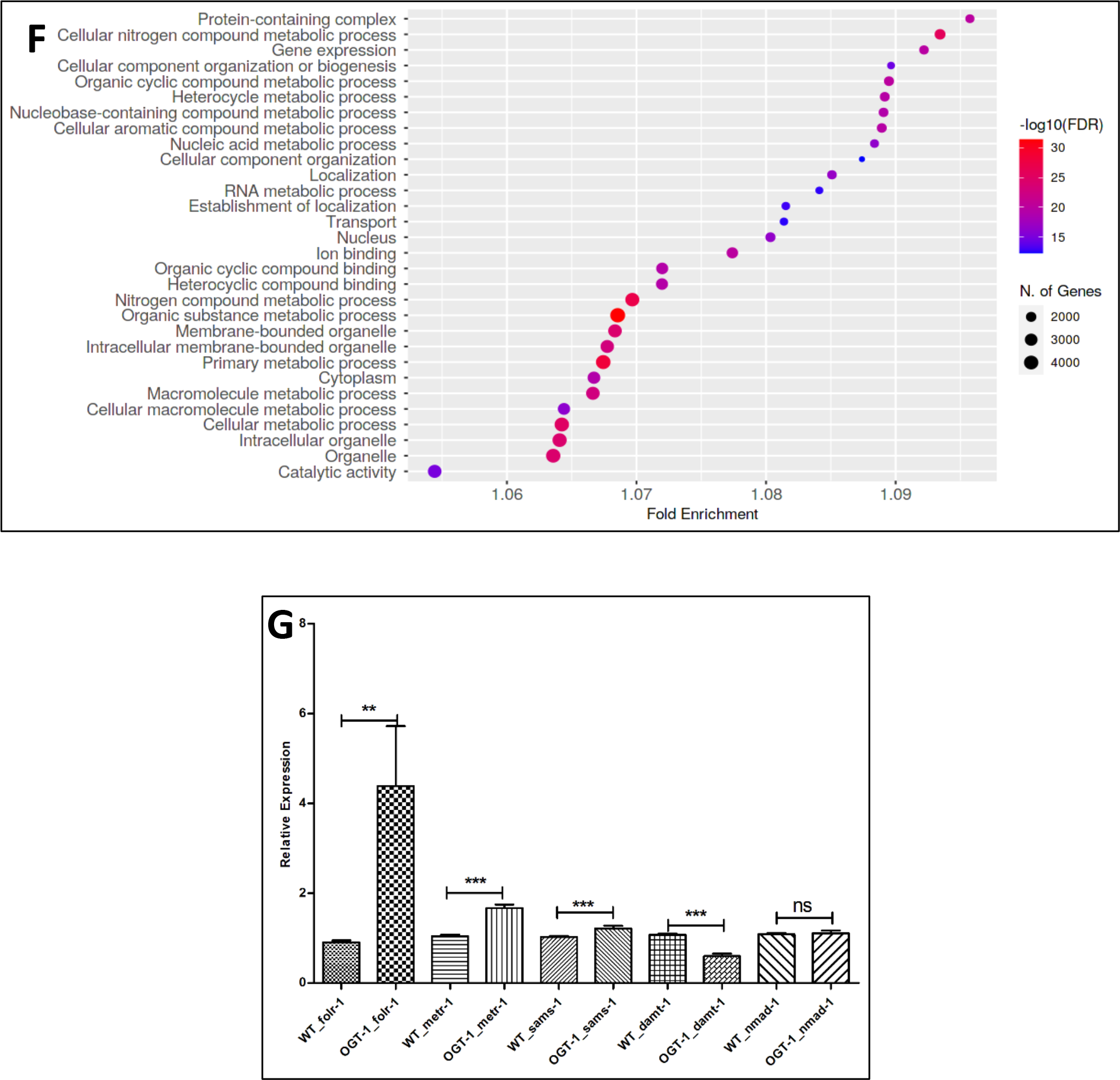
RNAseq data Analysis suggests important role of one carbon metabolism and related pathways in *ogt-1* mediated neuronal regeneration. **(A)** A scatter plot of differentially expressed genes (DEGs) identified in RNAseq between WT and *ogt-1* mutants. (B) Gene Ontology (GO) classification of DEGs in WT-vs-*ogt-1*. (C) KEGG pathway classification of differentially expressed genes (DEGs) in WT-vs-*ogt-1*. (D) KEGG pathway enrichment bubble plot of differentially expressed genes (DEGs). (E) Enrichment bubble plot of Gene Ontology molecular function analysis differentially expressed genes (DEGs). (F) Gene Ontology (GO) analysis of differentially expressed genes (DEGs) identified in neuron specific RNAseq between WT and *ogt-1* mutant (FDR0.1). (G) qRT-PCR of selected genes involved in OCM (*folr-1, metr-1 & sams-1*) and nucleic acid methyltransferases and demethylases (*damt-1 & nmad-2*). All data shown ±SEM, Student t-test; * p<0.05, **p<0.01, ***p<0.001.

To further investigate if OCM and its related pathways are influenced by *ogt-1* mutation specifically within neuronal cells, we performed RNAseq analysis in the RNA samples isolated from FACs (Fluorescence-activated cell sorting) sorted neuronal cells in WT and *ogt-1* worms (supp Fig. 2A). Neuron specific RNAseq analysis identified a significant number of differentially expressed genes (DEGs) (supp Fig. 2B, Table-S3). As with whole worm analysis, Gene Ontology (GO) pathway analysis of neuron specific DEGs identified metabolic processes such as cellular, macromolecule, nitrogen compound, nucleic acid metabolism *etc.* (Fig. 2F, Table-S3). GO analysis of twofold up regulated genes revealed neuron specific pathways as anticipated (neuronal perception, chemical and olfactory perception, synapses *etc*) along with carbohydrate and polysaccharide metabolic pathways (supp Fig. 2C, Table-S3), while twofold down regulated genes included biological processes like meiosis, mitosis, gamete/ germ cell production and maturation, reproduction, cell cycle, nuclear division and embryonic developments *etc.* which are expected to be down regulated in the neuronal tissue (supp Fig. 2D, Table-S3). Our top 50 up and down regulated genes (supp fig. 2E and 2F) include important genes regulated by *daf-2* and *daf-16* which have been reported to play critical role in adult neuron function and regeneration (Kaletsky, Lakhina et al. 2016). In addition, other important genes involve in metabolism, epigenetic modification and ATP metabolism are also enriched. Employing whole animal qRT-PCR, we further confirmed that *folr-1, metr-1, sams-1,* important genes for OCM, were significantly up regulated in the *ogt-1* background compared to WT (Fig. 2G). While DNA methyltransferase (*damt-1*) was significantly down regulated and DNA demethylases (*nmad-1*) was unchanged (Fig. 2G), Further bioinformatic analysis of neuron specific DEGs using the Functional Annotation Tool “DAVID Bioinformatics Resources” revealed enrichment of metabolic pathways such as glycolysis, lipid metabolism along with serine synthesis pathway (SSP), OCM, amino acid, nucleotide, and nitrogen compound metabolism *etc.* (supp Fig. 3A). While biosynthesis of cofactor analysis specified enrichment of Folate, Methionine and SAM metabolism cycles, glutathione metabolism and ATP synthesis pathways (supp Fig. 3B). Taken together, the results of our unbiased high throughput gene expression analysis strongly indicate the involvement of OCM and its offshoot pathways in the increased neuronal regeneration in *ogt-1* mutant animals.

**Fig.3. Functional.**
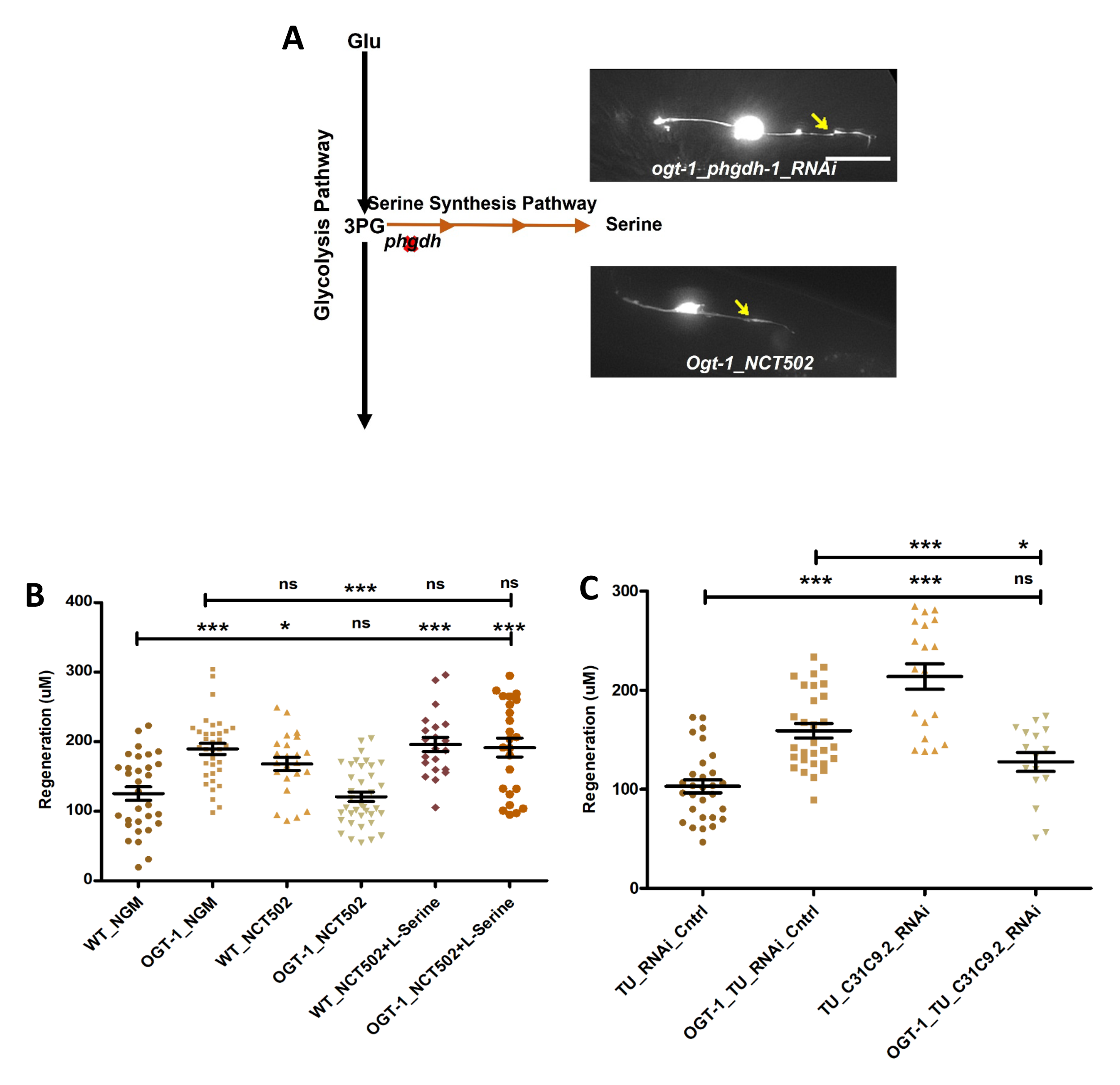
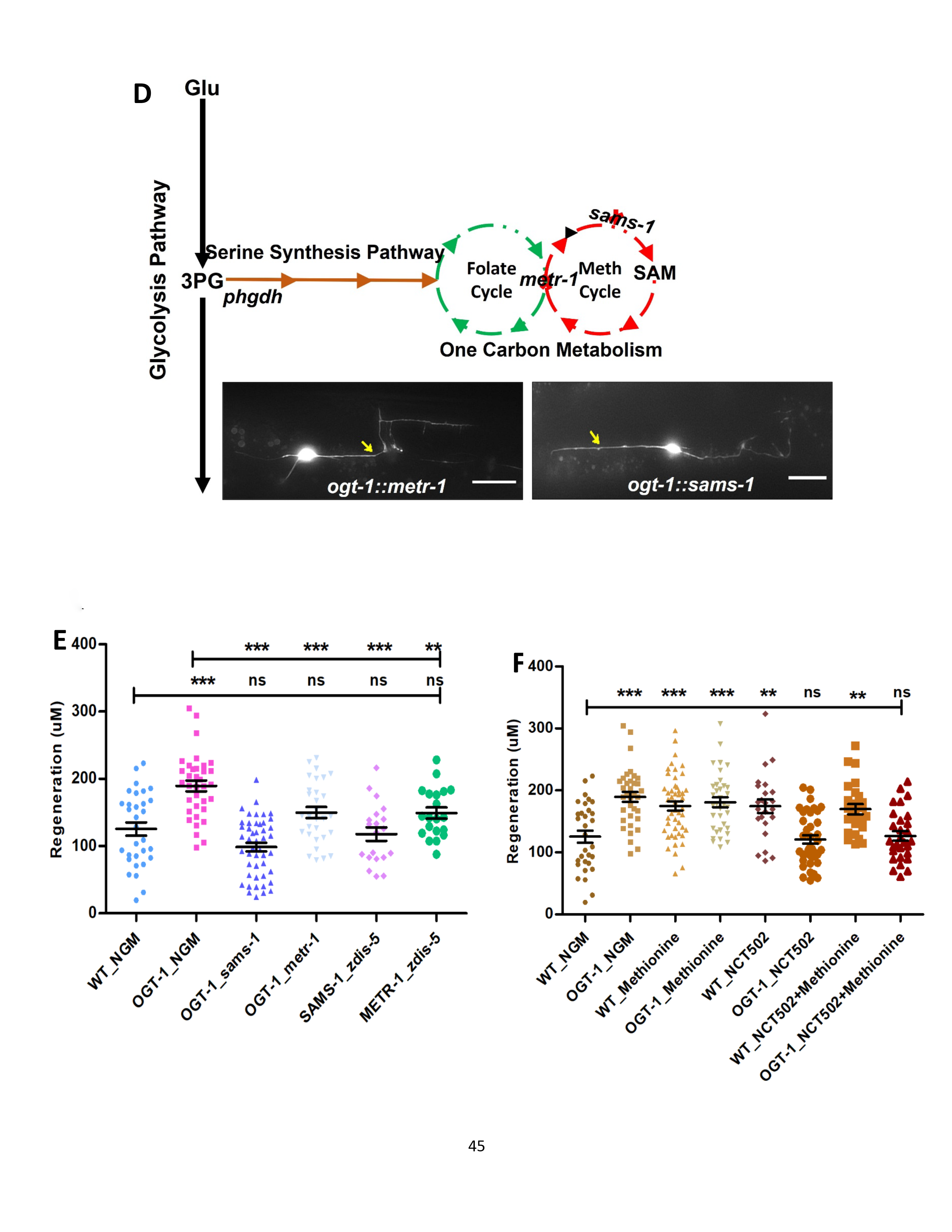
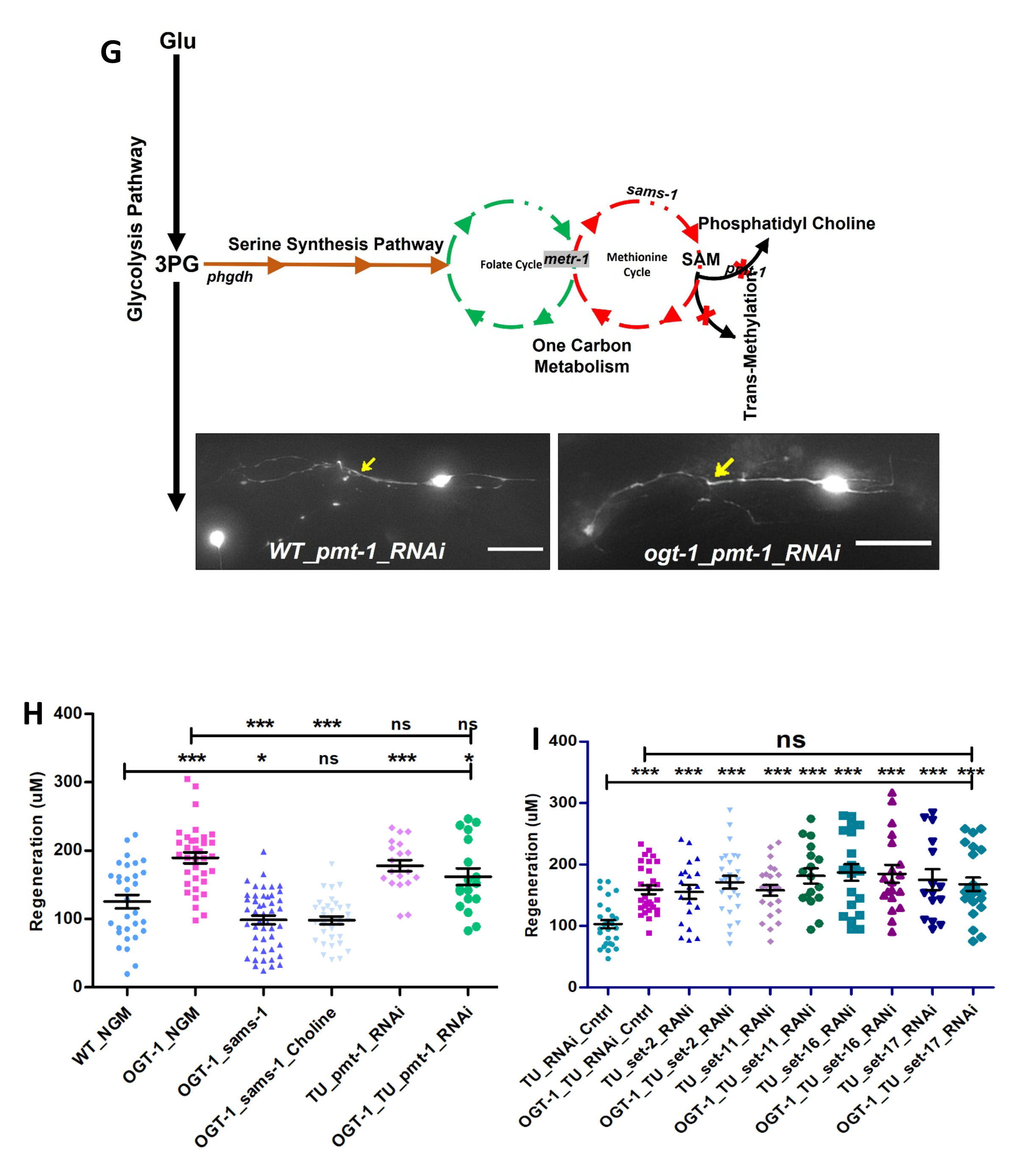
one carbon metabolism (OCM) and serine synthesis pathway (SSP) are essential for neuronal regeneration in *ogt-1* worms. **(A)** Schematic representation showing Glycolysis and the Serine Synthesis Pathway (SSP), along with representative images at 24 h neuron regeneration in conditions blocking the SSP in *ogt-1* mutants using either neuron specific RNAi or NCT502 drug (arrow indicates the point of injury). **(B)** Effect of NCT502 drug and supplementation of serine on WT and *ogt-1* mutant 24 h neuronal regeneration. **(C)** Effect of neuron specific RNAi against *C31C9.2* (ortholog of human *PHGDH* gene) on WT and *ogt-1* mutant neuronal regeneration. **(D)** Schematic representation of the metabolic link between glycolysis and OCM via SSP, along with images of 24 h neuron regeneration with OCM gene mutations. **(E)** Effects of *metr-1* and *sams-1* mutations on enhanced regeneration in *ogt-1* worms. **(F)** Effects of methionine supplementation on regeneration in WT, *ogt-1* animals, and on the *phgdh-1* inhibitor drug NCT502. **(G)** Schematic representation of OCM metabolite SAM usage in lipogenesis and transmethylation, along with images of neuron regeneration when they are blocked. **(H)** 24 h neuron regeneration with choline supplementation and neuron specific RNAi against *pmt-1*. **(I)** 24 h neuron regeneration when blocking methyltransferases by neuron specific RNAi (*set-2, set-11, set-16,* and *set-17*) in WT and *ogt-1* animals. scale bar = ∼10uM, all data shown in ±SEM, One Way ANOVA *pValue <0.05, **pValue <0.01, ***pValue <0.001.

### 3. Functional One-carbon Metabolism (OCM) and Serine Synthesis Pathway (SSP) are indispensable for enhanced regeneration in *ogt-1* animals

Following the result of our gene expression analysis we sought to functionally validate the importance of the OCM and related pathways in neuronal regeneration in *ogt-1* worms. We first focused on the serine synthesis pathway (SSP) as it metabolically connects glycolysis with OCM (Fig. 3A) (Yu, Wang et al. 2019). NCT502 (MCE HY-117240) is a chemical agent reported to inhibit the mammalian phosphoglycerate dehydrogenase (PHGDH) enzyme, which catalyzes the first and rate limiting step of serine biosynthesis (Tabatabaie, Klomp et al. 2010, Zogg 2014, Pacold, Brimacombe et al. 2016). Applied to *C. elegans*, NCT502 abrogated the effect of *ogt-1* mutation on neuronal regeneration but significantly increased the regeneration in WT worms (Fig. 3B, Table-S4). In addition, we observe that supplementation of L-serine, the final product of SSP, which feeds in to OCM, rescued the abrogative effect of NCT502 in *ogt-1* (Fig. 3B, Table-S4). Previously we found that AKT kinase, *akt-1,* activity, plays an important role in *ogt-1* regeneration, *akt-1* mutation blocked the enhanced regeneration of *ogt-1,* while gain of function *akt-1 (++)* phenocopied *ogt-1* effect (Taub, Awal et al. 2018). Interestingly, NCT502 blocked the enhanced regeneration *in ogt-1 (-); akt-1 (++)* worms (supp fig. 4A, Table-S6) and serine supplementation rescued the enhanced regeneration that is eliminated in *akt-1(-)*;*ogt-1(-)* worms (supp fig. 4A, Table-S6). Since NCT502 has not been earlier reported to be used in *C. elegans*, we also tested the effects of blocking SSP using RNAi gene knockdown. In concordance to NCT502 treatment, neuron specific RNAi against C31C9.2 (*phgdh-1*), the *C. elegans* ortholog of human PHGDH and target of NCT502, abrogated the effects of *ogt-1* mediated regeneration, and significantly increased the regeneration in WT worms even beyond that of *ogt-1* worms (Fig. 3C, Table-S4). Interestingly, systemic RNAi knockdown against C31C9.2 (*phgdh-1*), that is ineffective in neurons, did not alter regeneration levels in *ogt-1* animals suggesting a neuron specific mechanism. However, it did significantly increase regeneration in WT worms (Supp fig. 4B, Table-S6). We further measured *pyk-1* activity in WT worms and found that it was significantly enhanced by NCT502 treatment (supp fig. 4C) suggesting increased glycolytic activity upon blocking the SSP. Interestingly, we observed equally enhanced *pyk-1* activity in *ogt-1* worms with NCT502 treatment (supp fig. 4D). These results demonstrate the importance of the SSP pathway in *ogt-1* mediated enhanced neuron regeneration but suggest that in wild-type animals the reverse may be true and blocking SSP becomes beneficial.

**Fig. 4.**
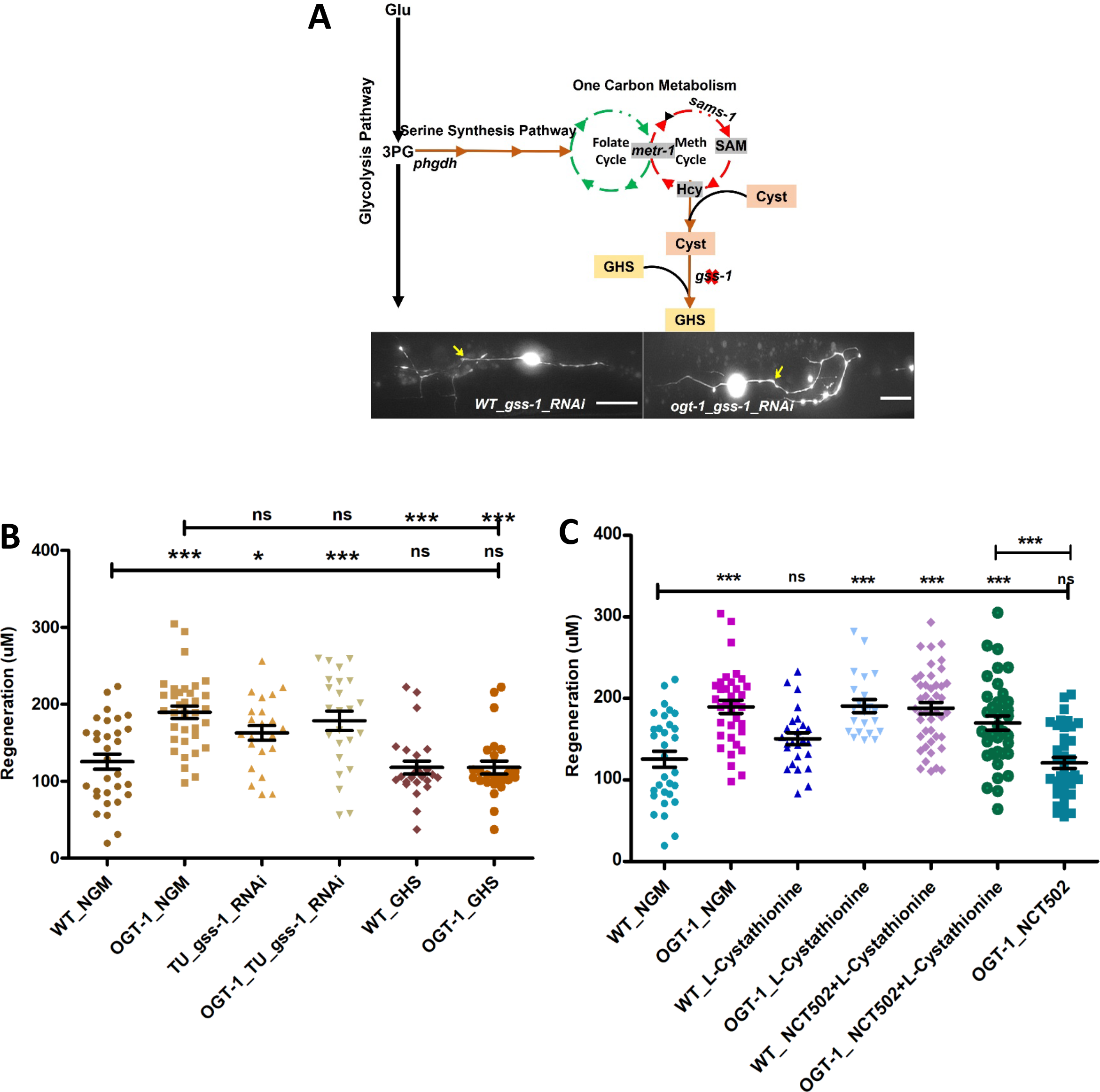
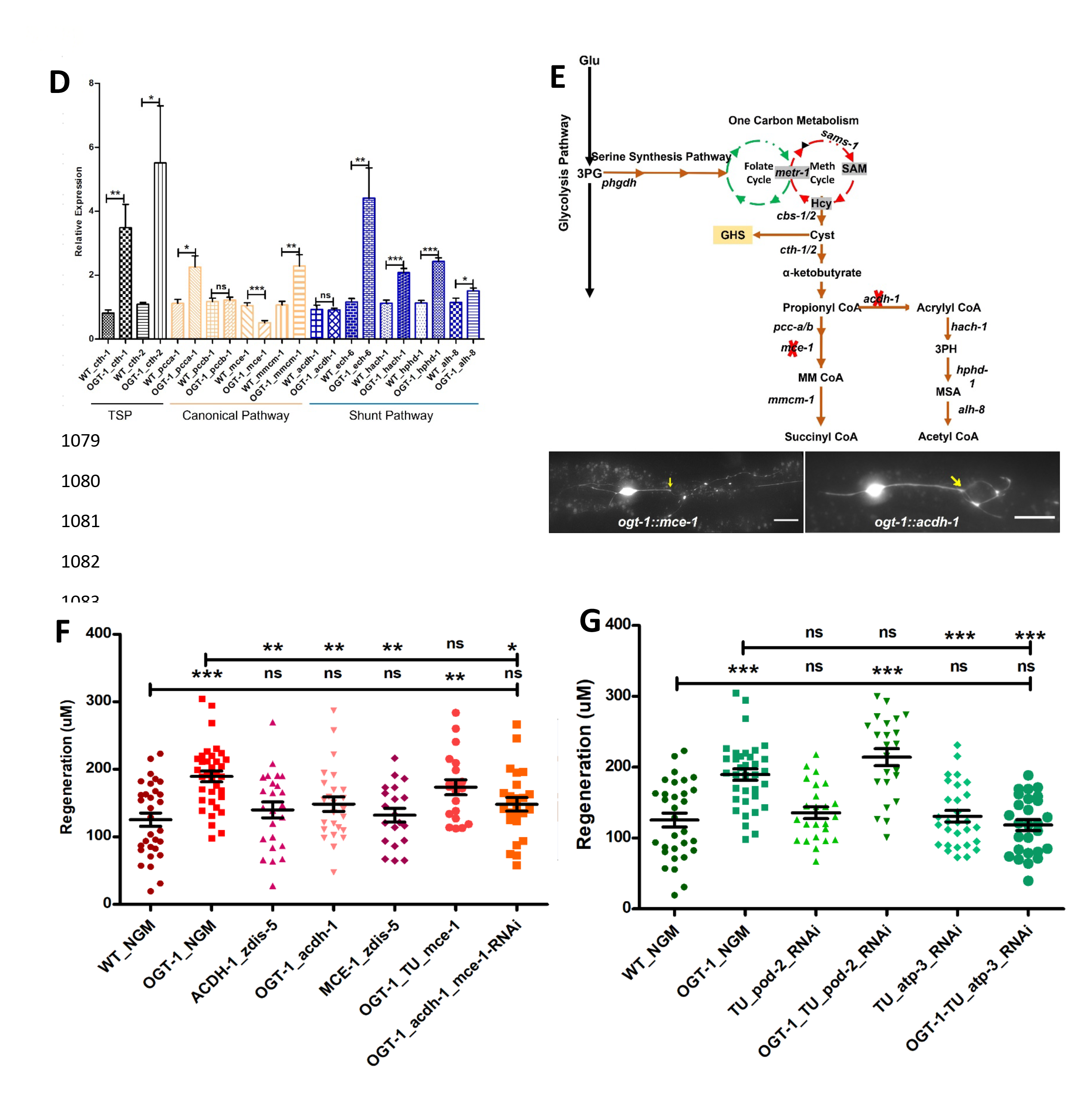
The transsulfuration pathway (TSP) leading to Acetyl-CoA production mediates enhanced regeneration in *ogt-1* animals: **(A)** Schematic representation of the transsulfuration pathway (TSP) branch of OCM, along with supplementation with TSP metabolites L-cystathionine, Glutathione and neuron specific RNAi against Glutathione synthetase (*gss-*1) with its effect on 24 h neuron regenerating neuron images (arrow indicates the point of injury). **(B)** Effects of GHS supplementation and neuronal RNAi knockdown against *gss-1* on neuronal regeneration in WT and *ogt-1* worms. **(C)** Effects of L-cystathionine supplementation on neuronal regeneration in WT and *ogt-1* worms, with or without SSP blocking by NCT502. **(D)** qRT-PCR of selected genes involved in transsulfuration (*cth-1* & *cth-2*), as well as the related downstream vitamin B12 dependent canonical pathways (*pcca-1, pccb-1, mce-1 & mmcm-1*) and the vitamin B12 independent Shunt pathway (*acdh-1, ech-6, hach-1, hphd-1 & alh-8*). **(E)** Schematic representation of the transsulfuration pathway (TSP) metabolites L-Cystathionine metabolism in to succinyl-CoA and Acetyl-CoA and genes involved with indicated mutant used in the study, along with representative regenerating neuron image (arrow indicates the point of injury). **(F)** Effect of *acdh-1* and *mce-1* mutation in WT and *ogt-1* background on neuronal regeneration. **(G)** Effect of blocking lipid synthesis from acetyl CoA and ATP production on regeneration in WT and *ogt-1*. Scale bar = ∼10uM, all data shown in ±SEM, analytical methods student t-test and One Way ANOVA were used *pValue <0.05, **pValue <0.01, ***pValue <0.001.l

To test the importance of OCM in *ogt-1* mediated regeneration directly, we tested mutations of methionine synthase (*metr-1*), an ortholog of the human MTR gene and s-adenosyl methionine synthetase-1 (*sams-1*), an ortholog of human MAT1A and MAT2A genes, in the *ogt-1* background (*ogt-1::metr-1* and *ogt-1::sams-1*) (Fig. 3D). Both mutations abrogated the enhanced regeneration in *ogt-1* animals but had no significant effect on WT regeneration (Fig. 3E, Table-S4). Methionine is an important metabolite of the OCM cycle and its supplementation increases OCM flux (Miousse, Pathak et al. 2017, Sanderson, Gao et al. 2019, Ligthart-Melis, Engelen et al. 2020). Methionine supplementation significantly increased the regeneration in WT worms but had no additional effect on *ogt-1* worms (Fig. 3F, Table-S4, and supp fig. 4E, Table-S6). Nor did it alter the effects of blocking the SSP in either WT or *ogt-1* animals (Fig. 3F, Table-S4) which may be in part due to the requirement of serine for normal OCM progression (Yang and Vousden 2016, Clare, Brassington et al. 2019, Geeraerts, Heylen et al. 2021). S-adenosyl Methionine (SAM), a product of SAMS-1 and an important metabolite of OCM, mediates numerous cellular processes including several biosynthetic, post-translational modifications and epigenetic modifications of histones and nucleic acids for regulation of gene expression and metabolism, including glycolysis (Ducker and Rabinowitz 2017, Clare, Brassington et al. 2019). It participates in the Kennedy pathway to synthesize lipid (Phosphatidyl Choline) an important component of the cellular membrane (Fig. 3G) (Walker 2017). Phosphatidyl Choline can alternatively be synthesized from choline. However we found that choline supplementation in *ogt-1::sams-1* dual mutant failed to rescue the effects of *sams-1* mutation (Fig. 3H, Table-S4). Furthermore, neuron specific RNAi against Phosphoethanolamine Methyl Transferase (*pmt-1*), involved in Phosphatidyl Choline biosynthesis from SAM, did not reduce *ogt-1* mediated regeneration, although it did enhance the regeneration in WT worms (Fig. 3H, Table-S4). SAM also acts as a methyl doner for transmethylation reactions including histone modification. To test if the epigenetic modification of histones by histone methyltransferases play any role in *ogt-1* enhanced regeneration, we knocked down several reported H3K4 methyltransferase with known effects on H3K4 methylation and/or neuronal regeneration including *set-2, set-11, set-16* and *set-17* (Walker, Jacobs et al. 2011, Wilson, Giono et al. 2020). Knocking down these methyltransferases had no significant effect on *ogt-1* mediated enhanced regeneration but significantly increased regeneration in WT worms (Fig. 3I, Table-S4). RNAseq analysis also showed that DNA methylases (*damt-1*) and demethylase (*nmad-1*) as well as *pmt-1/pmt-2*, required for Phosphatidyl Choline synthesis from SAM, were all relatively downregulated while OCM genes were relatively upregulated in neuronal tissue in *ogt-1* animals (supp fig. 4F). Thus, while the functional OCM pathway mediated by MERT-1 and SAMS-1 is essential for *ogt-1* mediated enhanced regeneration, these results suggest that it does not act through either lipogenesis or transmethylation pathways involved in epigenetic regulation.

### 4. The transsulfuration pathway (TSP), an offshoot of OCM, is critical for enhanced neuronal regeneration in *ogt-1* animals

Our gene expression analysis revealed that OCM related pathways such as glutathione and SAM metabolism are highly altered in *ogt-1* worms. We therefore tested the importance of the transsulfuration pathway in *ogt-1* mediated regeneration (fig. 4A). The transsulfuration pathway involves cysteine and cystathionine metabolism that is utilized in glutathione synthesis important for oxidative stress maintenance in neurons (Vitvitsky, Thomas et al. 2006, Sbodio, Snyder et al. 2019). Performing neuron specific RNAi against Glutathione Synthetase (*gss-1*), an ortholog of human glutathione synthetase (GSS), we detected no effect on the enhanced regeneration in the *ogt-1* mutant background but significantly increased regeneration in WT (fig. 4B, Table-S5). In a complimentary manor, supplementation with L-Glutathione (GHS) significantly decreased regeneration in *ogt-1* worms but had no effect on WT worms (fig. 4B, Table-S5). By contrast, supplementation with L-cystathionine had no detectable effect on regeneration in *ogt-1* worms or WT (fig. 4C, Table-S5) but rescued the effect of blocking SSP with NCT502 in *ogt-1* worms (fig. 4C, Table-S5). These observations suggest that while the transsulfuration pathway is functionally involved in *ogt-1* mediated enhanced regeneration it is not through glutathione synthesis.

Cystathionine can be further metabolized in to succinyl-CoA or acetyl-CoA through either the vitamin B12 dependent canonical pathway or the vitamin B12 independent shunt pathways respectively (Watson, Olin-Sandoval et al. 2016, Giese, Walker et al. 2020). Succinyl-CoA or acetyl-CoA can be further used for different metabolic processes or can enter the Krebs Cycle to produce ATP. Our neuronal cell specific RNAseq analysis revealed that genes involved in OCM (*metr-1, sams-1, folr-1, mthf-1 etc.*), transsulfuration (*cth-1*) (supp fig. 4F, 5A), and the downstream vitamin B12 independent shunt pathway (*acdh-1, ech-6, hach-1, hphd-1 & alh-8*) were relatively upregulated (supp fig. 5A) in *ogt-1* animals, while genes involved in the vitamin B12 dependent canonical pathway (*pcca-1, pccb-1, mce-1 & mmcm-1*) were down regulated (supp fig. 5A). Likewise, performing qRT-PCR analysis against genes in these pathways, we found that genes involved in TSP (*cth-1, cht-2*) and in the vitamin B12 independent shunt pathway showed unidirectional upregulated expression in *ogt-1* (fig. 4D), while genes involved in the canonical vitamin B12 dependent pathway showed no clear trend in differential expression (fig. 4D).

**Fig. 5.**
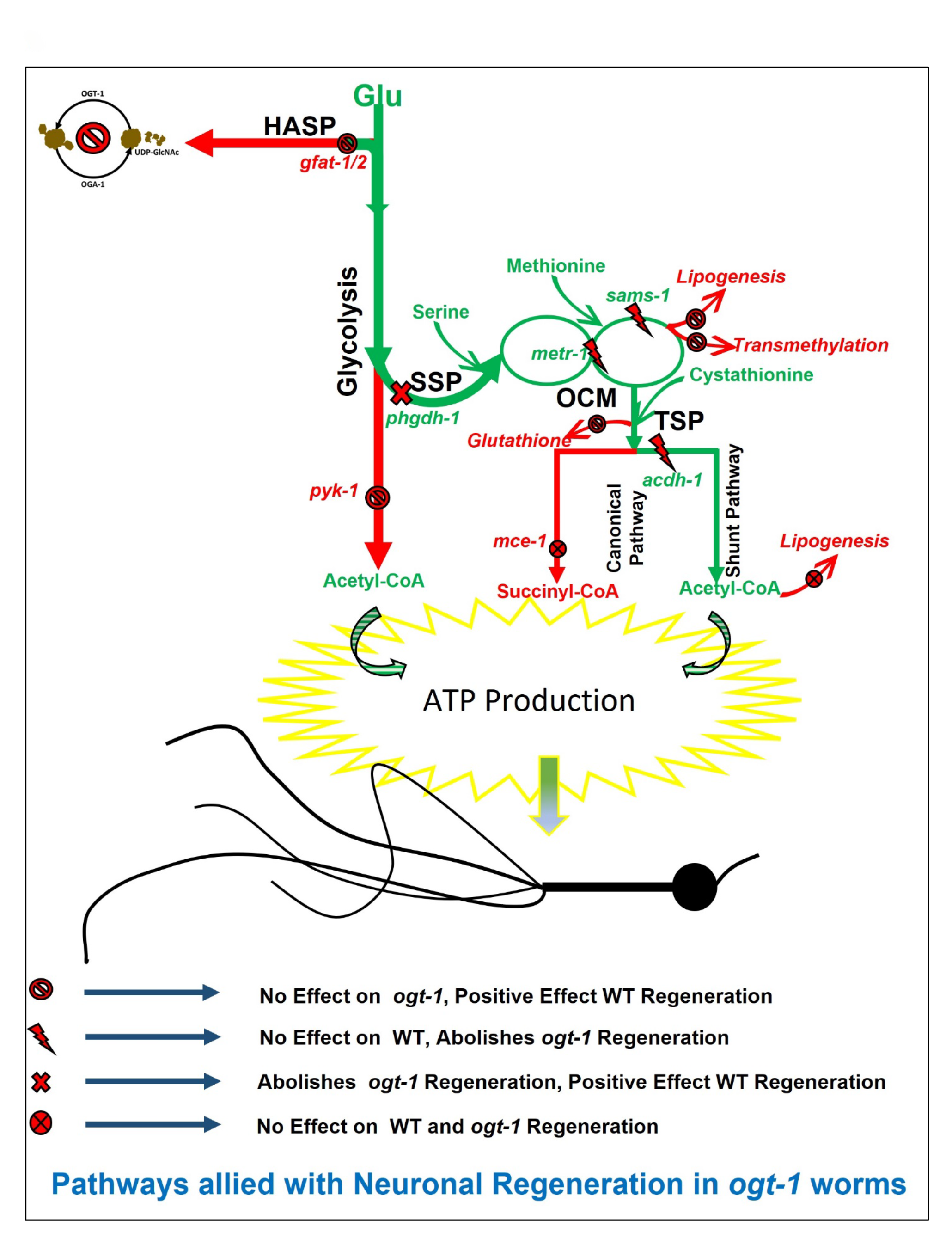
The metabolic pathway for enhanced neuronal regeneration in *ogt-1* animals: A detailed schematic of the metabolic pathway for the enhanced regeneration in *ogt-1* animals with the tested genes, metabolite supplementations and pharmacological treatments indicated. As highlighted in green, *ogt-1* mutations divert metabolic flux from enhanced glycolysis to OCM *via* the SPP, driving metabolites in the transsulfuration pathway (TSP) to support enhanced regeneration *via* the vitamin B12 independent shunt pathway. Dispensable metabolic branches are shown in red.

To test the role of cystathionine metabolism through shunt and canonical pathways directly in neuronal regeneration, we generated double mutants with acyl-CoA dehydrogenase (*acdh-1)* that mediates the vitamin B12 independent shunt pathway, *ogt-1::acdh-1,* and methylmalonyl-CoA epimerase (*mce-1*) that mediates the vitamin B12 dependent canonical pathway *ogt-1::mce-1* (fig. 4E). The *mce-1* mutation had no effect on regeneration in either WT or the *ogt-1* background (*ogt-1::mce-1*) (fig. 4F, Table-S5). However, while *acdh-1* mutation had no effect on WT regeneration, it selectively eliminated the enhance regeneration of the *ogt-1* background (fig. 4F, Table-S5). These results were recapitulated using neuron specific RNAi knockdown against *acdh-1* and *mce-1* in the *ogt-1* background (supp fig. 5B, Table-S6). Neuron specific RNAi against *mce-1* in the *ogt-1::acdh-1* double mutant had no observable effect (fig. 4F, Table-S5). The *acdh-1* mediated shunt pathway is involved in the production of acetyl CoA from L-Cystathionine which can be further used for several processes including lipid synthesis and/or ATP production. Thus, we tested if lipid synthesis plays a role by neuron specific RNAi against *pod-2* (acetyl-CoA carboxylase), an ortholog of human ACACA (acetyl-CoA carboxylase alpha), that is important for lipid synthesis from acetyl CoA, but found it had no effect on either WT or *ogt-1* regeneration (fig. 4G, Table-S5). In contrast, the enhanced regeneration in *ogt-1* worms was clearly blocked by neuron specific RNAi against *atp-3* RNAi that reduces cellular ATP production, (as described above earlier fig. 1C and fig. 4G, Table-S5). In combination with gene expression analysis, these results further define the pathway of *ogt-1* regeneration to specifically involve acetyl CoA production by cystathionine metabolism through the vitamin B12 independent shunt pathway.

## Discussion

In order to initiate and sustain the energetically demanding growth state required for effective regeneration there must be sufficient modulation of the underlying molecular and metabolic processes within the damaged neuron (He and Jin 2016). Numerous studies have focused on the molecular and genetic mechanisms involved in axonal regeneration (Sun, Shay et al. 2014, Chisholm, Hutter et al. 2016, Chung, Awal et al. 2016). Yet the role of metabolic function in neuron regeneration is relatively less explored, despite its clear role in determining regenerative capacity (Taub, Awal et al. 2018, Li, Sami et al. 2020, Yang, Wang et al. 2020). Previously, our group demonstrated that genetically altered O-GlcNAc levels can substantially enhance neuronal regeneration through modulation of the neuronal metabolic response (Taub, Awal et al. 2018). Exploiting the genetic and optical accessibility of *C. elegans*, we demonstrated that a reduction of O-GlcNAc levels (*ogt-1* mutation), a proxy for metabolic deficit, resulted in enhanced glycolysis that supports increased regenerative capacity (Taub, Awal et al. 2018) and (Fig. 1A). Disruption of glycolysis (genetic or pharmacological) selectively eliminates the enhanced regeneration of the *ogt-1* mutant (Taub, Awal et al. 2018). Glycolysis is a key energy source for neurons, particularly under energy-limiting conditions (Jang, Nelson et al. 2016) and in developing neurons that foster high axonal growth rates (Han, Baig et al. 2016, Zheng, Boyer et al. 2016, Han, Xie et al. 2020). We have further verified this by neuron specific RNAi knockdown of *atp-3*, which significantly reduces cellular ATP levels (Soto, Rivera et al. 2020) and blocks the enhance regeneration in *ogt-1* animals (fig 1C). While our previous work established that neuronal glycolysis is a key component of enhanced axonal regeneration following injury in *ogt-1* worms (Taub, Awal et al. 2018), key questions remained as to what specific metabolic pathways, within the enhanced glycolysis background, are amended and involved to support regeneration.

Our results indicate that a complex metabolic pathway beyond that of canonical glycolysis is involved in the enhanced regeneration in *ogt-1* animals (fig 5). In our earlier study, we demonstrated the importance of early glycolytic enzymes (*pfk-1.1*, and *pgk-3*) in the *ogt-1* effect (Taub, Awal et al. 2018). However, we found here that this does not extend to the complete glycolytic pathway as neuron specific disruption of pyruvate kinase (*pyk-1*), which catalyzes the final step of glycolysis to produce pyruvate, had no effect on regeneration in *ogt-1* (Fig. 1C). This is in accordance with the reported effects of O-GlcNAcylation on these enzymes. High O-GlcNAcylation decreases *pfk-1.1* function (Bacigalupa, Bhadiadra et al. 2018). Despite the fact that high O-GlcNAcylation also destabilizes the pyruvate kinase, PKM1/2, complex (Wang, Liu et al. 2017), reports show that inhibition of *ogt-1* results in low pyruvate kinase expression and cellular activity (Yu, Teoh et al. 2019). The *ogt-1* mutation, which reduces O-GlcNAcylation, is therefor expected to increase *pfk-1.1,* and reduce *pyk-1*, activity respectively, which agrees with their measured importance in *ogt-1* neuron regeneration.

As these results indicate that the increased regeneration in *ogt-1* mutants does not entail direct ATP production in the TCA cycle of canonical glycolysis, we further adopted an unbiased approach performing genome wide gene expression analysis to identify additional pathways involved. Through Gene ontology (GO) and Kegg pathway classification analysis of RNAseq data from wild-type and *ogt-1* mutant animals we identified several metabolic pathways altered in both whole animals and FACs sorted neuron samples (fig. 2A and supp Fig. 2B). In addition to numerous genes and cellular processes with known roles in regeneration such as amino acid, nucleotide metabolism, lipid synthesis, methylation and glycolysis (Ducker and Rabinowitz 2017, Clare, Brassington et al. 2019), our analysis further identified metabolic processes including glutathione and s-adenosyl methionine (SAM) metabolism, energy metabolism and ATP synthesis that were significantly enriched in the *ogt-1* background. This pathway enrichment analysis indicates the involvement of One Carbon Metabolism (OCM) and its associated pathways in enhanced regeneration in *ogt-1* animals (fig2 and supp fig. 3). These results were further confirmed *via* specific gene expression analysis using qRT-PCR (fig 2G and fig. 4D) and indicate the importance of OCM and the Transsulfuration Pathway (TSP) as key metabolic pathways altered by the *ogt-1* mutation (Fig. 2D-E).

OCM is involved in a wide array of cellular processes including biosynthesis (purines and thymidine), amino acid homeostasis (glycine, serine, and methionine), epigenetic maintenance (nucleic acid and histone methylation), and redox defense (Ducker and Rabinowitz 2017). Enhanced glycolysis drives OCM through the Serine Synthesis Pathway (SSP) (Locasale 2013, Yu, Wang et al. 2019) that is known to be involved in several neuronal conditions including, neuronal growth, neural tube defect and Alzheimer’s disease (Coppedè 2010, Bonvento and Bolaños 2021, Lionaki, Ploumi et al. 2022). Through a combination of genetic manipulation, pharmacological treatment, and metabolic supplementation in our *C. elegans* neuronal regeneration assays, we have determined the specific metabolic pathway by which OCM contributes to the enhanced regeneration in the *ogt-1* mutant. The complete pathway is illustrated in green in Figure 5. We found that metabolic flux from the early steps of glycolysis is diverted to OCM through SSP, which is in agreement with earlier reports where enhanced glycolysis diverts metabolic flux towards OCM through SSP (Yu, Wang et al. 2019). This was most dramatically illustrated by the reduction in regeneration from pharmacological, or genetic, disruption of *phgdh-1* (*C31C9.2,* ortholog of human *PHGDH*), a key element of the SSP. The role of the SSP was further confirm by serine supplementation in the *akt-1* and *ogt-1* double mutant (*ogt-1;akt-1*), which restored the enhanced *ogt-1* regeneration blocked by the *akt-1* mutation (supp Fig. 4A). These results are in agreement with earlier metabolomic findings that enhanced glycolysis (Yu, Wang et al. 2019) and/or knock down of PMK1/2 (mammalian ortholog of *pyk-1*) diverts metabolic flux toward serine synthesis pathway to sustain cellular metabolic requirements (Yu, Teoh et al. 2019).

Although OCM is involved in both lipogenesis and DNA transmethylation (Kersten 2001, Yu, Wang et al. 2019) that could potentially play significant roles in increasing neuron regeneration (Iskandar, Rizk et al. 2010), we found that the regeneration effects of *ogt-1* were primarily dependent on L-cystathionine metabolism *via* the downstream TSP (fig. 4C). The TSP is influenced by OCM and its metabolites and has been reported to play an important role in neurodegenerative diseases and ATP production (Giese, Walker et al. 2020, Lam, Kervin et al. 2021). We found that cystathionine supplementation rescued the prohibitory effects of blocking the SSP pathway in the *ogt-1* background (fig. 4C). Testing branches of the TSP, we found that only the vitamin B12 independent shunt pathway was required, *via* Acyl CoA dehydrogenase (*acdh-1)*, for *ogt-1* mediated enhanced regeneration. The shunt pathway generates Acetyl-CoA that will drive ATP production through the Kreb’s cycle ultimately bringing the metabolic consequences of *ogt-1* back to cellular energy production and utilization as we demonstrated in Taub et al. Though we observed a significant decrease in ATP levels (Fig. 1F & supp Fig. 1C) and no difference in ATP utilization (supp Fig. 1D) in *ogt-1* animals, these observations maybe due to the fact that the measurements were either in whole worm or in nonneuronal tissues rather than neuron specific. Indeed, the down regulation of *pyk-1* from *ogt-1* inhibition has been associated with total reduced ATP levels previously (Dey, Son et al. 2019). Regardless, our work here has now deciphered the specific metabolic pathway through which the enhanced regenerative effect of *ogt-1* occurs.

While the *ogt-1* mutant diverts metabolic flux through a specific pathway to support and sustain enhance regeneration, we also discovered several additional conditions where restriction or diversion of metabolic flux in wild-type animals has similar beneficial effects. For instance, the HBS pathway nominally shunts off ∼5% of glycolytic flux (Marshall, Bacote et al. 1991, Bond and Hanover 2015). We found that blocking the HBS pathway through RNAi against *gfat-1* and *gfat-2* (Yi, Clark et al. 2012, Jóźwiak, Forma et al. 2014, Kim, Nakayama et al. 2018), appears to divert metabolic flux towards glycolysis and results in enhanced regeneration in WT animals similar to that of *ogt-1* (Fig. 1B). Likewise, *pyk-1* knockdown increases regeneration in WT and is known to divert metabolic flux toward the SSP (Yu, Teoh et al. 2019). Within OCM, we found that transmethylation pathways required for epigenetic modifications and phospholipid synthesis were not essential for the enhanced regeneration in *ogt-1* animals but that blocking histone methyl transferases in WT animals (Fig. 3I) increased regeneration. In addition, supplementation in wild type with the metabolite, L-methionine (product of *metr-1*), which increases OCM, phenocopied the enhance regeneration of the *ogt-1* mutant (fig 3F) as did blocking neuronal glutathione synthesis within the TSP (*gss-1* RNAi) (Fig 4B). While in the above instances restriction or enhancement of specific metabolic steps could be augmenting the same pathway utilized in *ogt-1* regeneration, in other cases clearly alternative pathways are at work. For example, pharmacologically (NCT502 treatment) or genetically (*phgdh-1* knock down) blocking SSP which restricts the *ogt-1* regeneration pathway effectively increases regeneration in WT. This effect is possibly due to increased metabolic flux through glycolysis, as we observed increased activity of *pyk-1* after NCT502 treatment (supp fig. 4C). Likewise, we had previously found that mutation of the O-GlcNAcase, *oga-1*, which increases O-GlcNAc levels, also increased neuron regeneration in *C. elegans,* but did so through an independent pathway of enhanced mitochondrial stress response (Taub, Awal et al. 2018).

Thus, within the complex web of cellular metabolism and energy production there appears to be numerous pathways for metabolite utilization that are beneficial for neuron regeneration. Here, employing genetic tools, we have defined the specific metabolic pathways (glycolysis, SSP, OCM and TSP) through which the *ogt-1* mutation diverts metabolic flux to increase neuronal regeneration. It is important to emphasize the accessibility of these metabolic effects to pharmacological treatment and/or metabolite supplement. For example, we previously demonstrated increased regeneration in wild-type animals with glucose supplementation (Taub, Awal et al. 2018). Here we find similar effects with L-methionine supplementation or treatment with the SSP blocking agent NCT502 in wild-type animals. Nutrient supplements and metabolic drug targets have been employed in neurotherapeutic treatments and prevention in numerous contexts including neuronal developmental defects (Greene, Leung et al. 2017, Businaro, Vauzour et al. 2021, Wu, Gao et al. 2022) and age associated neurodegenerative diseases (Stempler, Yizhak et al. 2014, Businaro, Vauzour et al. 2021). Our work demonstrates the necessities of OCM, SSP and TSP metabolic pathways and their interaction in the increased regenerative capacity of a damaged neuron in *ogt-1* animals and further highlights the distinct possibilities for such metabolic targets in the treatment of neuronal injury.

## Methods Details

### KEY RESOURCES TABLE

| REAGENT | RESOURCE SOURCE | IDENTIFIER |
| --- | --- | --- |
| **Bacterial Strains** |  |  |
| E. coli OP50 | Caenorhabditis Genetics Centre | OP50 |
| Vidal and Ahringer RNAi Libraries in HT115 (D3) E. coli | Source Bioscience | N/A |
| **Chemicals and Pharmacological Reagents** |  |  |
| L-methionine | Fisher Scientific | Cat#: AC166160025 CAS: 63-68-3 |
| L-Cystathionine | Sigma | Cat#: C7505-10MG |
| L-Methionine | Sigma | Cat#: M-9625 |
| L-Serine | Sigma | Cat#: S-4500 |
| Choline Chloride | Millipore Sigma | Cat#: C7017-5G, CAS:67-48-1 |
| Sodium Chloride (NaCl) | Fisher Bioreagents | Cat#: BP358-1 CAS:7647-14-5 |
| β-mercaptoethanol | Fisher Scientific | Cat#: AC125470100 CAS: 60-24-2 |
| Trizol | ThermoFisher Scientific | Cat#: 15596018 |
| PowerUp SYBR Green Master Mix | Applied Biosystems | Cat#: A25741 |
| Agarose | Fisher Bioreagents | Cat#: BP160-500, CAS: 9012-36-6 |
| Pronase | Sigma- Aldrich | SKU# 10165921001 |
| DMSO (Dimethyl Sulfoxide) | ThermoFisher Scientific | Cat#: 85190, Cas:67-68-5 |
| Water, Molecular Grade, Sterile, DEPC Free | Fisher Scientific | Cat#: R91450001G, CAS: 7732-18-5 |
| NCT502 | MedChemExpress (MCE) | Cat#: HY-117240 |
| Polybead polystyrene | Polysciences | Cat#08691-10 |
| **Commercial kit used** |  |  |
| ATP Assay Kit (Colorometric/ Fluorometric) | Abcam | Ab83355 |
| Pyrophosphate Assay Kit (Fluorometric) | Abcam | Ab112155 |
| Pyruvate Kinase (PK) Assay Kit (Colorimetric) | Abcam | Ab83432 |
| BCA Protein Quantification Kit | Abcam | Ab102536 |
| RNAeasy columns | QIAGEN | Cat#74034 |
| Direct-zol RNA Miniprep Plus Kit | Zymo Research | Cat# R2070 |

| Experimental models: Organisms/strains |  |  |
| --- | --- | --- |
| Worm | Strain | Source |
| WT (zdis5 pmec-4::GFP) | SK4005 | Taub *et. al.* |
| *ogt-1(ok1474)*_zdis-5 pmec-4::GFP | NA | Taub *et. al.* |
| *ogt-1(ok1474)* | RB1342 | Taub *et. al.* |
| TU3568 (sid-1(pk3321) him-5(e1490) V; lin-15B(n744) X; uls71[(pCFJ90) pmyo-2::mCherry + pmec-18::sid-1]) | NA | Taub *et. al.* |
| *ogt-1(ok1474)*_TU3568 | NA | Taub *et. al.* |
| *sams-1(ok3033)* | RB2240 | CGC |
| *sams-1*_zdis-5 | NA | in this study |
| *ogt-1;sams-1*_zdis-5 | NA | in this study |
| *metr-1(R03D7.1(ok521))* | RB755 | CGC |
| *metr-1*_zdis-5 | NA | in this study |
| *ogt-1;metr-1*_zdis-5 | NA | in this study |
| *acdh-1(ok1489)* | VC1011 | CGC |
| *acdh-1*_zdis-5 | NA | in this study |
| *ogt-1;acdh-1*_zdis-5 | NA | in this study |
| *mce-1(D2030.5(ok243))* | RB512 | CGC |
| *mce-1*_zdis-5 | NA | in this study |
| *mce-1*_TU3568_zdis-5 | NA | in this study |
| *ogt-1;mce-1*_TU3568_zdis-5 | NA | in this study |
| *ogt-1;akt-1(mg144)* | CG122 | Taub *et. al.* |
| *ogt-1;akt-1(ok525)* | CG125 | Taub *et. al.* |
| irls158 (normal ATP sensor, CAmA) | MS2495 | Soto and Rivera *et. al.* |
| *ogt-1; CAmA* | NA | In this study |

### REAGENTS AND RESOURCES

Further information and requests for resources, data and reagents should be directed to and will be fulfilled by the Lead Contact, Christopher V. Gabel.

### EXPERIMENTAL MODEL AND SUBJECT DETAILS

All *C. elegans* strains were cultured and maintained at 20^0^C on Nematode Growth Media (NGM) agar plates seeded with OP50 *E. coli*, unless otherwise noted. Strains were obtained from the Caenorhabditis Genetics Consortium (CGC at the University of Minnesota). To visualize the mechanosensory neurons, strains were crossed either into SK4005 (zdis5 [pmec4::GFP]) or *ogt*-1::zdis-5. Strains used are listed in detail in the experimental model table above. All strains generated by crossing were confirmed by genotyped using primers recommended by the CGC.

#### Laser Axotomy

*In vivo* Laser Axotomy was performed with a Ti:Sapphire infrared laser system (Mantis PulseSwitch Laser, Coherent Inc), that generated a 1 mHz train of 100 fs pulses in the near infrared (800 nm), pulse energy of 15-30 nJ/pulse. Axotomy was performed on a Nikon Ti-2000 inverted fluorescent microscope with a Nikon 40X 1.4 NA objective. Neurons were imaged for axotomy and subsequent measurement of regeneration *via* standard wide-field fluorescence of gfp expressed in the targeted neuron. Day 1 adult *C. elegans* were mounted on 5-6% agarose pads and immobilized in a 3-5 μL slurry of polystyrene beads (Polysciences, Polybead Polystyrene, 0.05 μM microsphere, cat#08691-10) and NGM buffer (Kim, Sun et al. 2013). Axotomy consisted of 3-5 short laser exposures (0.25 s each) resulting in vaporization at the focal point and severing of the targeted axon. The Anterior Lateral Microtubule (ALM) neuron was injured with two targeted cuts. The first cut was made 20 μm from the cell soma and a second cut was made 40-50 μm from the cell soma, creating a 20-30 μm gap. Regeneration was reimaged with a Nikon 40X 1.4 NA objective 24 h after axotomy, or as otherwise indicated, by placing the animals on a 2% agarose pad with 5mM sodium azide. Regeneration lengths were measured by tracing along the new neuron outgrowth with ImageJ/FIJI.

#### Mechanosensory Neuron Specific RNAi Feeding

To evaluate the function of specific genes, RNAi gene knockdown was employed following protocols we used previously, Taub *et al*. These protocols where first confirmed by performing RNAi knockdown against GFP in mechanosensory neurons and visually confirming significant reduction in GFP expression. Both the Ahringer and Vidal bacterial RNAi libraries were employed (Kamath and Ahringer 2003, Rual, Ceron et al. 2004). Following standard protocols, bacteria colonies were streaked out on LB agar containing penicillin and grown at 37^0^C overnight. The next day, single colonies were selected and grown in 10 mL of LB with Ampicillin overnight at 37^0^C. From this subculture, 250 micro-liters (uL) was spread onto RNAi agar plates containing penicillin and 2mM IPTG. Plates were dried and incubated at room temperature for at least 48 hours before using them for worm culturing. For mechanosensory neuron specific RNAi gene knockdown, we employed the TU3568 (sid-1(pk3321) him-5(e1490) V; lin-15B(n744) X; uIs71 [(pCFJ90) pmyo-2::mCherry + pmec-18::sid-1]) background (Calixto, Chelur et al. 2010). This strain has RNAi sensitivity specifically in the mechanosensory neurons and is RNAi resistant in all other tissues. TU3568 was crossed into the *ogt-1* mutant background. Following protocols we established in Taub et al 2018, gravid adults were bleached, and embryos were allowed to hatch onto RNAi-bacteria plates. Once the F1 generation reached adulthood, 30-40 gravid adults were picked onto fresh RNAi-bacteria plates and allowed to lay eggs for 3-4 hours. The day 1 adults of the F2 generation were then used for Laser Axotomy and regeneration assays as described above. Animals were rescued on a fresh RNAi plate and cultured until imaging was performed.

#### Drug Treatments in *C. elegans*

For all chemical reagent and metabolite treatments, the compound was dissolved in NGM agar before being poured into plates. Animals were cultured on treated plates for their lifespan before and after axotomy. Choline 30mM (Sigma, Cat#: C7017-5G), (Ding, Smulan et al. 2015), L-Methionine 75 μM (Fisher Scientific, Cat#: AC166160025), 5 mM L-Serine (Sigma, cat# S-4500) (Liu, Janssens et al. 2019), L-Cystothionine 50 μM (Sigma, Cat#: C7017-5G, CAS:67-48-1) and L-Glutathione reduced 100 μM (Cat#: G4251-50G), were dissolved in molecular grade water (Fisher Scientific, Cat#: R91450001G, CAS: 7732-18-5) at required stock concentrations (Ellwood, Slade et al. 2022). The phgdh (C31C9.2) inhibitor N-(4,6-dimethylpyridin-2-yl)-4-[5-(trifluoromethyl)pyridin-2-yl]piperazine-1-carbothioamid (NCT502) (MedChemExpress, Cat#: HY-117240) was initially dissolved in DMSO and diluted in ddH2O to use at a concentration of 25 μM in NGM plates (Pacold, Brimacombe et al. 2016). (note: we have shown previously that DMSO does not affect regeneration (Taub, Awal et al. 2018).

#### qRT-PCR

To evaluate the expression levels of candidate genes in wildtype and *ogt-1* animals we performed qRT-PCR. Day 1 adult *C. elegans* were lysed in 0.5% SDS, 5% b-ME, 10 mM EDTA, 10 mM Tris-HCl pH 7.4, 0.5 mg/ml Proteinase K, then RNA was purified with Tri-Reagent (Sigma). DNAse I treatment (NEB M03035) of 2-3 ug RNA followed by cDNA conversion using High-Capacity cDNA Reverse Transcription Kit (Thermo Fisher Scientific cat#4368814). qRT-PCR was performed in biological triplicate with three technical triplicates for each condition using Real-Time PCR Quantstudio 12K Flex qPCR System and Fast SYBR Green Master Mix (Thermo Fisher, 4385617). Relative transcript abundance was determined by using the DDCt method and normalized to *act-1* mRNA expression levels as a control. Primers are listed in Table-S7.

#### Neuronal cell isolation from adult animals using FACs

To isolate neuronal cells from Day 1 adult worms we utilized the protocol developed and described earlier (Zhang, Banerjee et al. 2011, Kaletsky, Lakhina et al. 2016). In brief WT (*unc-119:*:GFP) and *ogt-1* (*ogt-1*::*unc-119*::GFP) worms expressing GFP in all neurons were generated by crossing WT or OGT-1 worms with otIs45 [*unc-119*::GFP]. Synchronized day 1 adult worms were washed (3X) with s-basal buffer to remove excess bacteria. The packed worm volume (250-350 μl) was washed twice with 500 μl lysis buffer (200 mM DTT, 0.25% SDS, 20 mM HEPES pH 8.0, 3% sucrose) and resuspended in 1,000 μl lysis buffer. Worms were incubated in lysis buffer with intermittent gentle tapping for 10 minutes at room temperature. The pellet was washed 6X with s-basal and resuspended in 20 mg/ml pronase solution from Streptomyces griseus (Sigma-Aldrich, SKU# 10165921001). Worms were incubated at room temperature (15-20 min) with periodic mechanical disruption by pipetting at every 2 min intervals. When most worm bodies were dissociated, leaving only small debris and eggs (as observed under a dissecting microscope), dissolved whole worm tissues were filtered to remove eggs and single cells were pelleted down at 4K RPM for 20 minutes at 4^0^C. The pellets were resuspended in ice-cold PBS buffer containing 2% fetal bovine serum (Gibco). The resulting dissociated cell suspension was subjected to Fluorescence-activated cell sorting (FACs) to isolate GFP labeled neurons (supp Fig. 2A).

#### Expression profiling by RNA-seq

Gene expression patterns in WT and *ogt-1* mutants were measured by RNAseq analysis from RNA extracted from both, day 1 adult, whole animal and FACs sorted neuronal cells. RNA from FACS-sorted neurons was extracted using the Direct-zol RNA Miniprep Plus Kit (Zymo Research, R2070). RNA from whole animals was extracted manually by lysing day 1 adult *C. elegans* in 0.5% SDS, 5% b-ME, 10 mM EDTA, 10 mM Tris-HCl pH 7.4, 0.5 mg/ml Proteinase K, then RNA was purified with Tri-Reagent (Sigma cat# T9424-25ML). Isolated RNA was purified by RNAeasy columns (QIAGEN, Cat#74034) and quality of RNA was evaluated with the 2100 bioanalyzer (Agilent) before library generation for the RNAseq experiments. RNA-seq experiments were not randomized, nor results blinded, as all analysis is fully automated and unbiased. For whole-worm and neuron-specific RNA sequencing of adult animals N = 2 biological replicates were used. No statistical methods were used to predetermine sample size (Kaletsky, Lakhina et al. 2016).

For whole body RNAseq analysis we acquired DNBseq RNA sequencing services from BGI Global (https://gtech.bgi.com/bgi/home). Total RNAseq and data analysis was performed by using BGI Global inhouse developed sequencing methods and data analysis. In brief, transcriptome libraries were generated using the library conversion kit before sequencing was performed on the DNBseq platform. For each library, 10 ng library was used to incorporate a 5′ phosphorylation, on the forward strand only, using polymerase chain reaction (PCR). Purified PCR product with 5′ phosphorylation was then denatured and mixed with an oligonucleotide ‘splint’ that is homologous to the P5 and P7 adapter regions of the library to generate a ssDNA circle. A DNA ligation step was then performed to create a complete ssDNA circle of the forward strand, followed by an exonuclease digestion step to remove single stranded non-circularized DNA molecules. Circular ssDNA molecules were then further subjected to Rolling Circle Amplification (RCA) to generate DNA Nanoballs (DNB) containing 300–500 copies of the libraries. Each DNB library was then drawn through a flow cell ready for sequencing using the DNBseq platform to generate 30 M clean reads per sample. FASTQ files were generated locally at sequencing performed by BGI. After data cleaning, processing includes removing adaptors, contamination, and low-quality reads. Bowtie2 was used to map the clean reads to the reference gene sequence (transcriptome), and then RSEM was used to calculate the gene expression level of each sample. The DEseq2 method was used to detect differentially expressed genes (DEGs).

For neuron specific RNAseq analysis we employed the Illumina NextSeq 2000 RNA sequencing services from “The Boston University Microarray & Sequencing Resource” (https://www.bumc.bu.edu/microarray/). RNA isolated from FACs sorted neuronal cells were subjected to quality control assessment using a bioanalyzer (Aligent). mRNA enrichment, library preparation and quality assessment were performed according to manufacturer protocols (Illumina). Sequencing was performed on the Illumina NextSeq 2000 System using the NextSeq 2000, P2 Reagent Kit (100 cycles) with sequencing read length 50x50 paired end. Sequencing data were assessed for the quality of each sample using **FastQC** (https://www.bioinformatics.babraham.ac.uk/projects/fastqc/), and **RSeQC** (https://rseqc.sourceforge.net/). Each sample was aligned to the genome using **STAR** (https://github.com/alexdobin/STAR), and **SAMtools** (https://samtools.sourceforge.net/) was used to count proper pairs of reads aligning to mitochondrial or ribosomal RNA. The subread package: high-performance read alignment, quantification and mutation discovery **featureCounts** (https://subread.sourceforge.net/) was used for alignment of proper read pairs unique to non-mitochondrial Ensemble Genes. As a control, all reads were also aligned to the GFP sequence, which indicated that all samples were GFP-positive as expected. To identify genes whose expression changes significantly between genotypes, a one-way analysis of variance (ANOVA) was performed using a likelihood ratio test to obtain a p value for each gene. Benjamini-Hochberg False Discovery Rate (FDR) correction was applied to obtain FDR-corrected p values (q values), which represent the probability that a given result is a false positive based on the overall distribution of p values. The FDR q value was also recomputed after removing genes that did not pass the “independent filtering” step in the DESeq2 package. Wald tests were then performed for each gene between experimental groups to obtain a test statistic and p value for each gene. FDR correction was then applied, across all genes for which a p value could be computed for all comparisons and across only those genes that passed expression filtering.

#### RNA Seq Bioinformatic Analysis

The following unbiased enrichment analysis was used to understand whether the differentially expressed gene list identified in the RNAseq data was significantly enriched in a pathway, molecular function, or particula biological process. **Gene Ontology (GO)** was employed to determine the molecular function, cellular component, and biological process of the differentially expressed genes. All differentially expressed genes where mapped to terms in the Gene Ontology database (http://www.geneontology.org/), the number of genes in each term calculated and a hypergeometric test applied to identify GO terms that are significantly enriched in candidate genes compared to the background of all genes in the species. In addition, we also utilized the online ShinyGO v0.741: Gene Ontology Enrichment Analysis (http://bioinformatics.sdstate.edu/go74/) to analyze neuronally enriched genes. **KEGG Pathway-based analysis** (Qvalue ≤ 0.05) was employed to determine the most important biochemical metabolic and signal transduction pathways significantly enriched in the differentially expressed genes. The differentially expressed gene list was further analyzed for functional annotation of enriched pathways using The **Database for Annotation, Visualization, and Integrated Discovery (DAVID)**. These tools are powered by the comprehensive DAVID Knowledgebase built upon the DAVID Gene concept which pulls together multiple sources of functional annotations. Using the recommended protocol for analysis in wizard tool of DAVID (https://david.ncifcrf.gov/tools.jsp) we analyzed the pathways and metabolites most affected in neurons of *ogt-1* mutants.

#### ATP Quantification via the FRET based ATP Sensor

We obtained the worm strain (MS2495) expressing the Fluorescence resonance energy transfer (FRET) based ATP sensor (novel**C**lover-**A**TP-**mA**pple fusion protein; **CAmA**) under the *pept-1* promoter expressed in the intestinal cells from Dr. Morris F Moduro lab (Soto, Rivera et al. 2020). Clover is a green fluorescent protein that is excited by blue light (480nm-510nm laser) and emits green light (511nm-530nm). mApple is a red fluorescent protein that is excited by green light (522nm-577nm) and emits red light (580nm-675nm). The *ogt-1* mutant was crossed with the ATP sensor strain (MS2495). The anterior gut of day1 adult worms (control and *ogt-1* mutant) was imaged to measure FRET fluorescence using a 63x objective on a confocal Zeiss LSM 880 microscope. Following established FRET imaging protocols, a mApple image was acquired first *via* direct excitation (561nm laser) and emission (594nm) to assess where the sensor protein was present and establish a baseline measurement. A second image was then obtained using a FRET filter set, i.e. excitation of Clover (488nm laser), producing green emission (522nm-577nm) that excites mApple which is detected as red emission (516nm) (FRETred). ImageJ was used to quantify the relative FRET pixel intensity (FRETred/baseline) within the region of interest.

#### ATP and Pyrophosphate (PPi) Quantification and *pyk-1* Activity Assay

Synchronized day 1 adult worms were collected in S-basal buffer and were washed 3x with s-basal and 1x in ATP assay buffer (Abcam, Ab83355), followed by sonication on ice in ATP assay buffer using a model 110V/T Ultrasonic Homogenizer for two cycles of 15 minutes. Sonicated samples were then centrifuged at 13,000 RPM for 15 minutes at 4^0^C. The supernatant was collected and moved to a fresh microcentrifuge tube and ATP quantitation was performed with the ATP Assay Kit (Colorometric/ Fluorometric) (Abcam, Ab83355) using a Tecan Infinite M1000 Pro Multi Microplate Reader. ATP was normalized to protein content measured with the BCA Protein Quantification Kit (Abcam, Ab102536). Triplicate technical replicates were performed for each sample; at least three biological samples were assayed for each condition reported. For *pyk-1* activity and pyrophosphate PPi quantification assays, animals were cultured as in ATP quantification assays and animals were sonicated on ice in respective assay buffers (*pky-1* or PPi assay buffer) and activity was recorded using a Tecan Infinite M1000 Pro Multi Microplate Reader. To normalize samples, the BCA Protein Quantification Kit (Abcam, Ab102536) was used.

#### Quantification and Statistical Analysis

Statistical analysis and graph generation was performed with Prism (Graph Pad). All data were compared with either WT, *ogt-1* mutant or RNAi control regeneration data. Data are shown as the mean with error bars representing the standard error of the mean. One-way ANOVA analysis with Dunnett’s and *post hoc* Bonferroni’s correction was employed for multiple comparisons. When only two groups of data were compared an unpaired t test was employed. In all cases, *p < 0.05 **p < 0.01, ***p < 0.001.

## SUPPLEMENTAL INFORMATION

Supplemental Information includes four figures and one table and can be found with this article online at

## ACKNOWLEDGMENTS

## Acknowledgements

We would like to thank Dr. Amy Walker, UMASS Worcester, MA, USA, Dr. Morris Maduro, UC Riverside, and The *C. elegans* Genetics Center provided many of the strains and Boston University Core (Dr. Tilton, Brian Richard (FACs); Dr. Yuriy Alekseyev (RNA sequencing and data analysis), Dr. Trinkaus-Randall (confocal Zeiss LSM 880 microscopy), and Dr. Lyn and Au, Matthew Bo (qRT-PCR) facilities for maintaining and making available different instrument for use. Dr. Walker provided *sams-1* worm, Dr. Maduro provided ATP expressing strain. Dr Danial Taub provided feedback and suggestions on the manuscript. Funding was provided by the Massachusetts Spinal Cord Injury Cure Research Program, INTF3110HH2191525007, from the Massachusetts Department of Public Health.

## AUTHOR CONTRIBUTIONS

D.K.Y. and C.V.G. conceived and designed experiments. D.K.Y. and C.V.G. performed all the experiments and aided in the analysis of data. A.S.C. aided in confocal imaging and analysis.

D.K.Y. and C.V.G. wrote the manuscript with input from all authors.

## Supplementary Figures

**Supplemental Figure 1.**
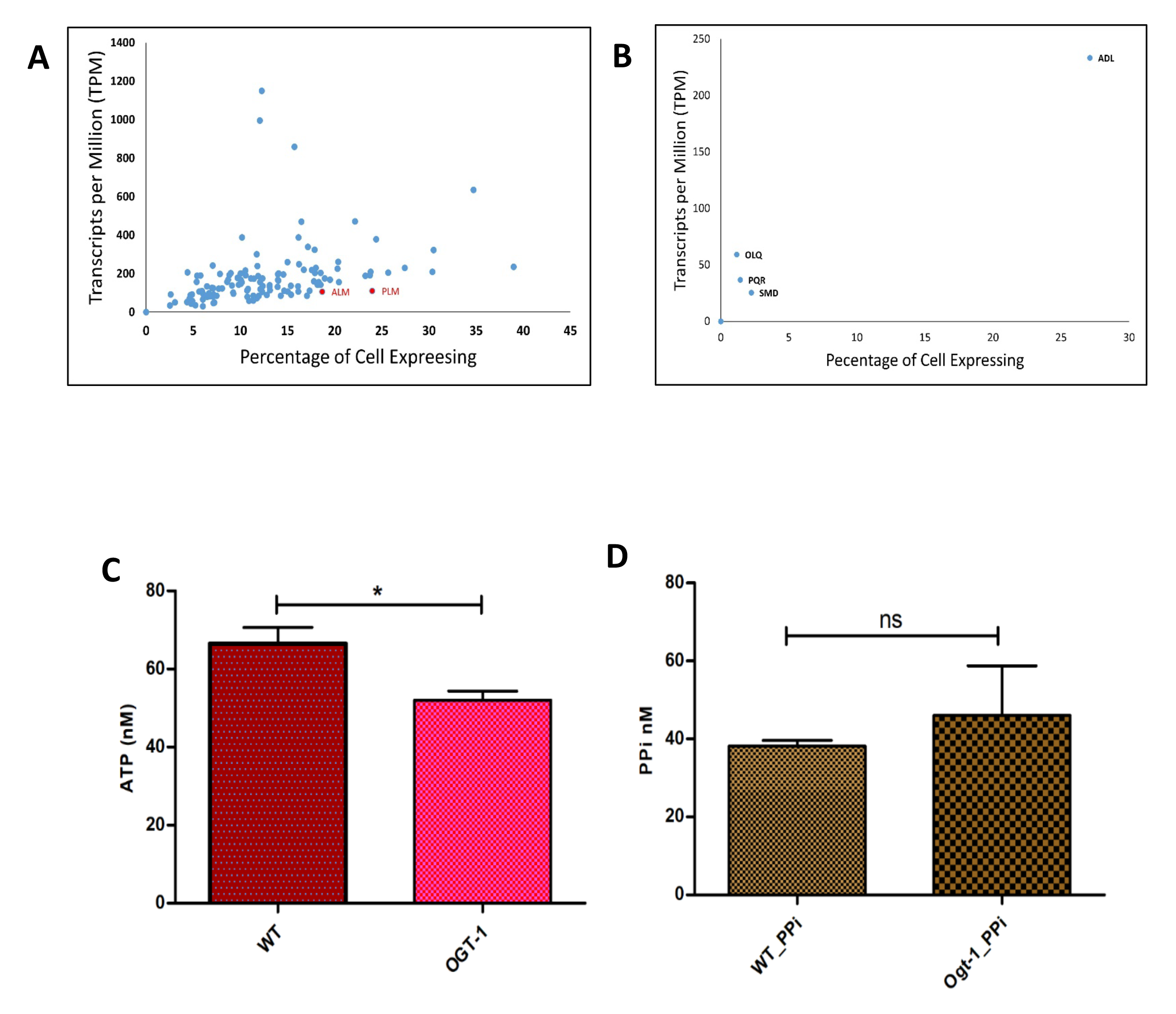
**(A)** *pyk-1* expression analysis in neuronal cell, single cell neuronal RNAseq data from worm base (https://wormbase.org/species/c_elegans/gene/WBGene00009126#0-9fce6b37d81-10) was used to generate the image. **(B)** *pyk-2* expression analysis in neuronal cell, as for *pyk-1.* **(C)** Relative amount of ATP measured using ATP Assay kit (Abcam, cat# Ab83355) in whole worm lysate. **(D)** Relative amount of Pyrophosphate (PPi) measured using Pyrophosphate Assay kit (Abcam, cat# Ab112155) in whole worm lysate. All data shown in ±SEM, analytical methods, student t-test and One Way ANOVA were used *pValue <0.05, **pValue <0.01, ***pValue <0.001.

**Supplemental Figure 2.**
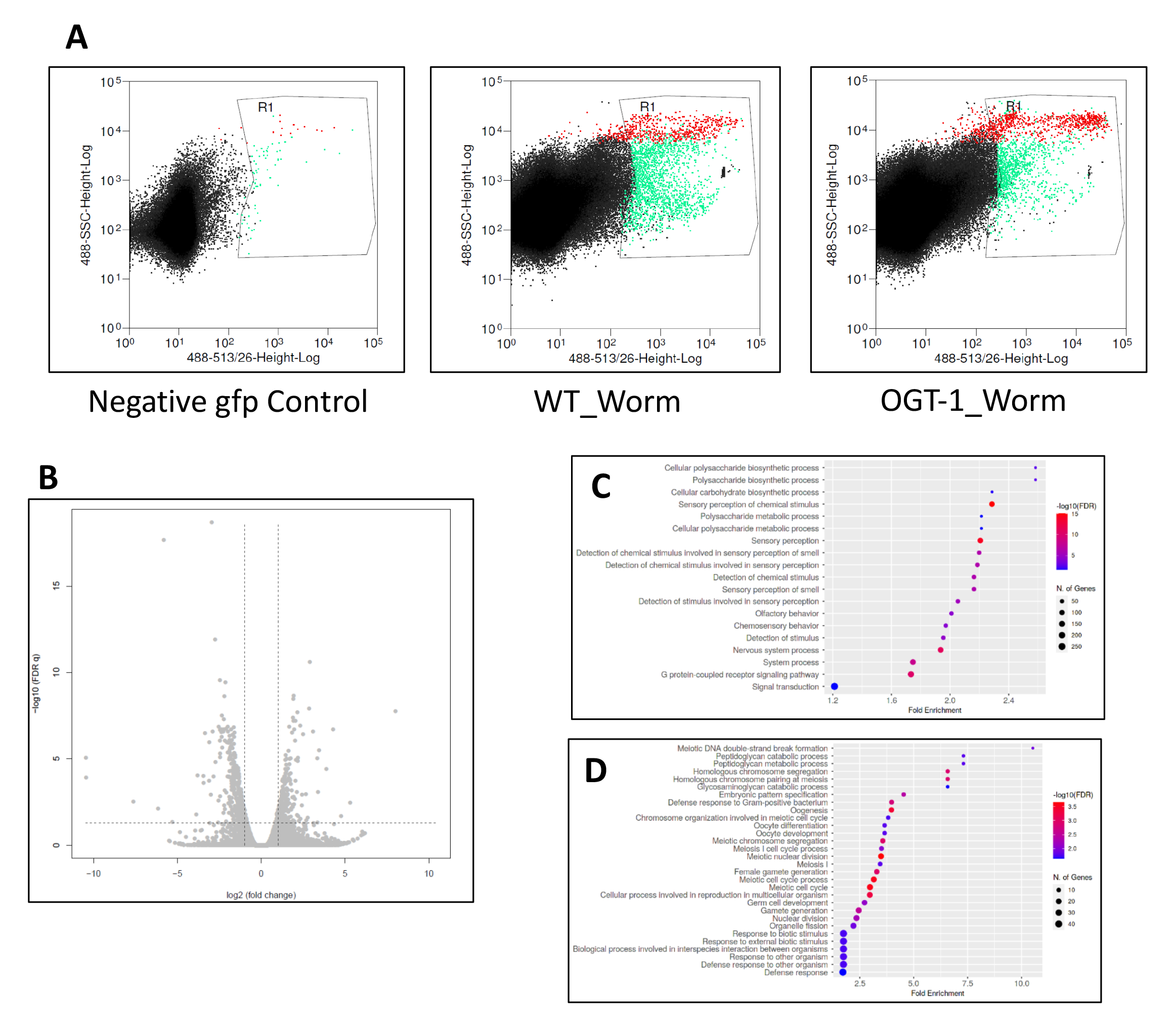
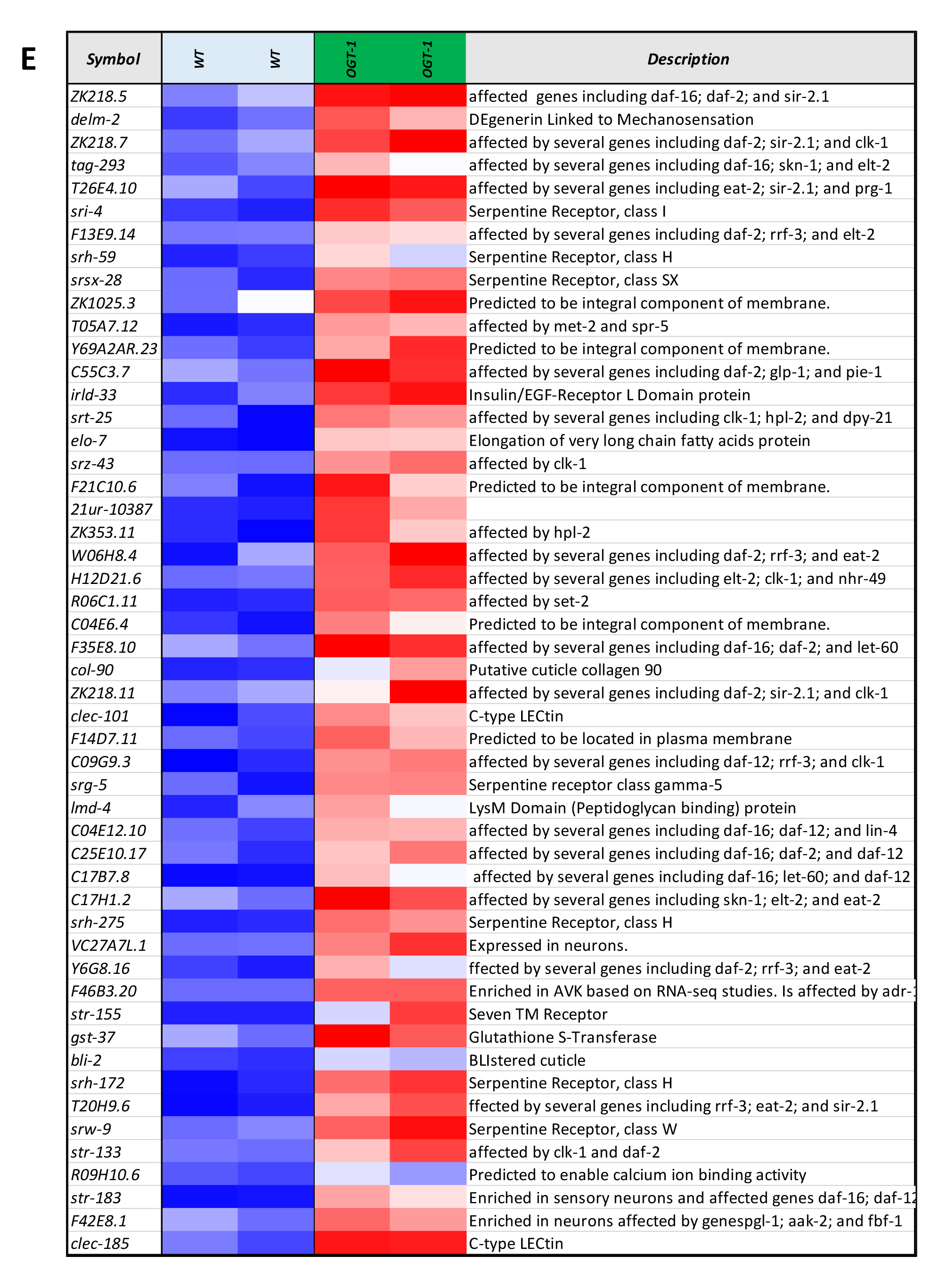
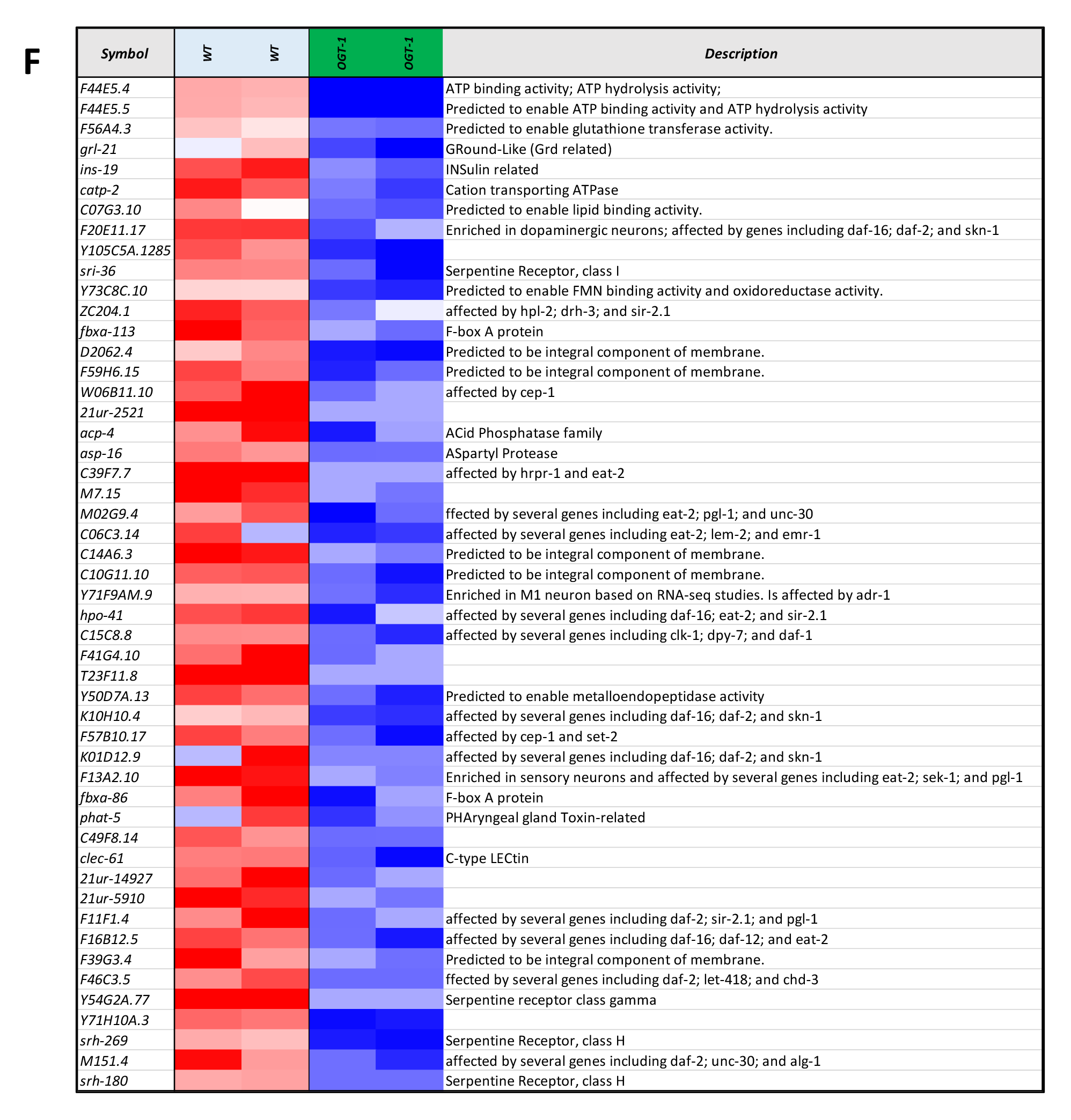
**(A)** Representative image of FACs sorting for GFP tagged neuronal cells used for RNA isolation and RNAseq analysis. GFP control (left), wild type (middle) and *ogt-1* mutant (right) worms, respectively. **(B)** Volcano plot for differentially expressed genes (DEGs) FDR0.05. **(C)** Gene Ontology (GO) analysis of 2fold up regulated DEGs in WT-vs-*ogt-1* (FDR0.1) **(D)** Gene Ontology (GO) analysis of 2fold down regulated DEGs in WT-vs-*ogt-1* (FDR0.1). **(E)** List of top 50 up regulated genes, and **(F)** top 50 down regulated genes and their function, in *ogt-1* animals, identified in neuron specific RNAseq.

**Supplemental Figure 3.**
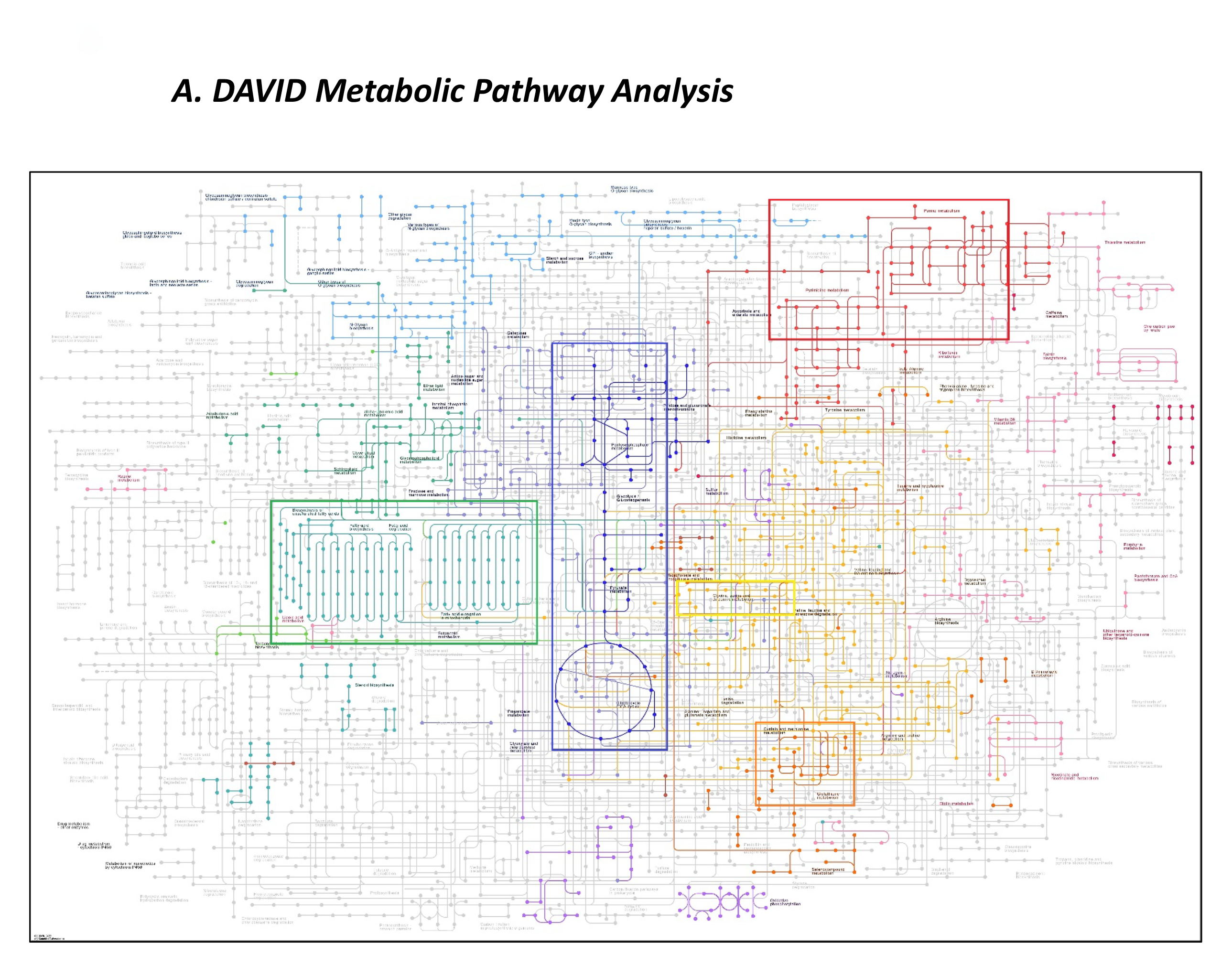
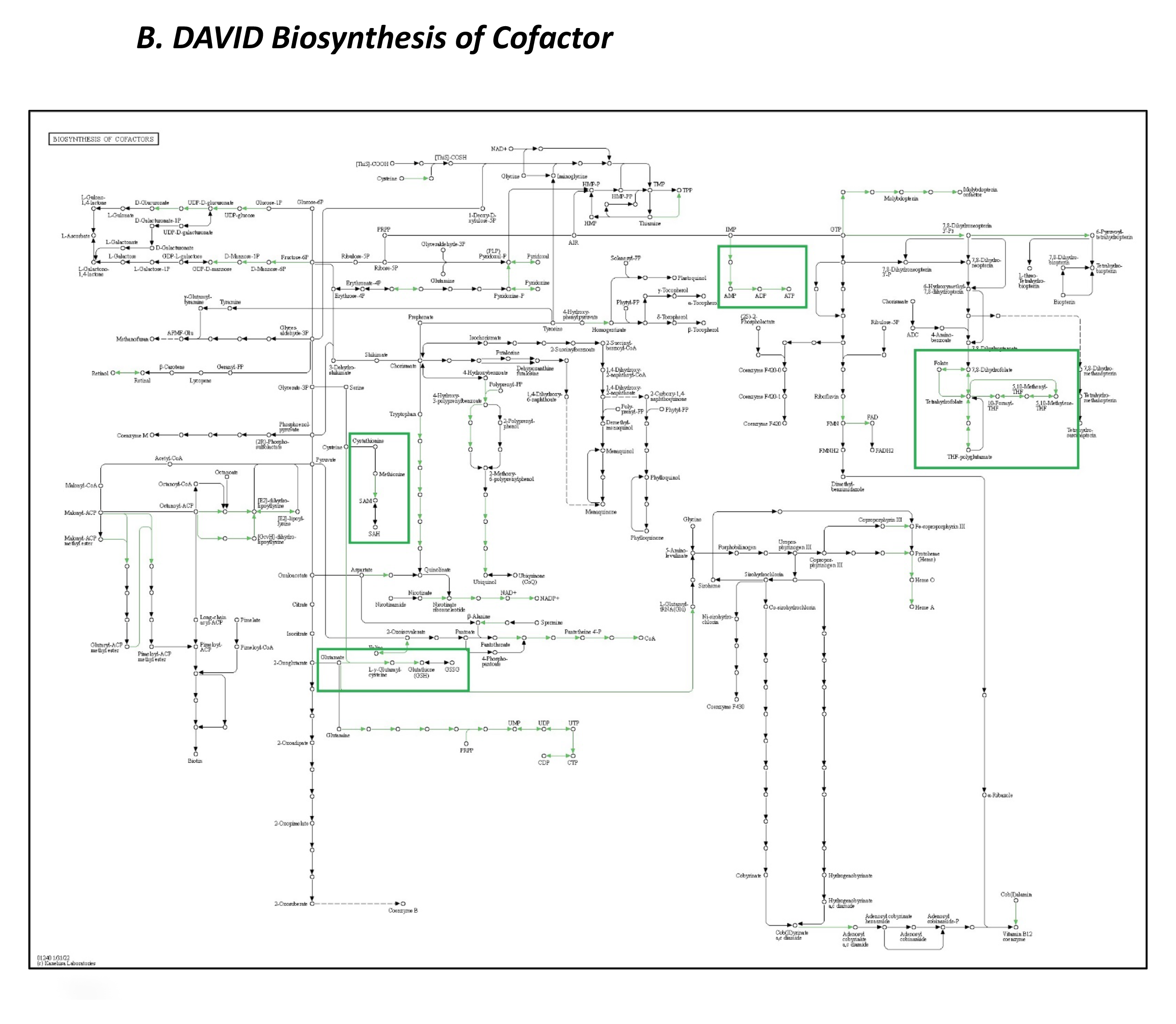
**(A)** Visualization of metabolic pathway enriched in differentially expressed genes (FDR0.1) identified in neuron specific RNAseq analysis using “DAVID Metabolic Pathway Analysis” tool. Top highlighted pathways are glycolysis (blue); lipid metabolism (green); nucleotide metabolism (red); serine synthesis pathway (light yellow) and one carbon metabolism and related pathways (dark yellow) respectively. **(B)** Pathway analysis of co-factor mediated biosynthesis of differentially expressed genes (FDR0.1) identified in neuron specific RNAseq analysis using the “DAVID Biosynthesis of Cofactors Analysis” tool. Most affected pathways (green highlighted) include those related with One Carbon Metabolism (folate, methionine and SAM metabolism); Transsulfuration pathway (Cystein & Glutathione metabolism) and ATP production.

**Supplemental Figure 4.**
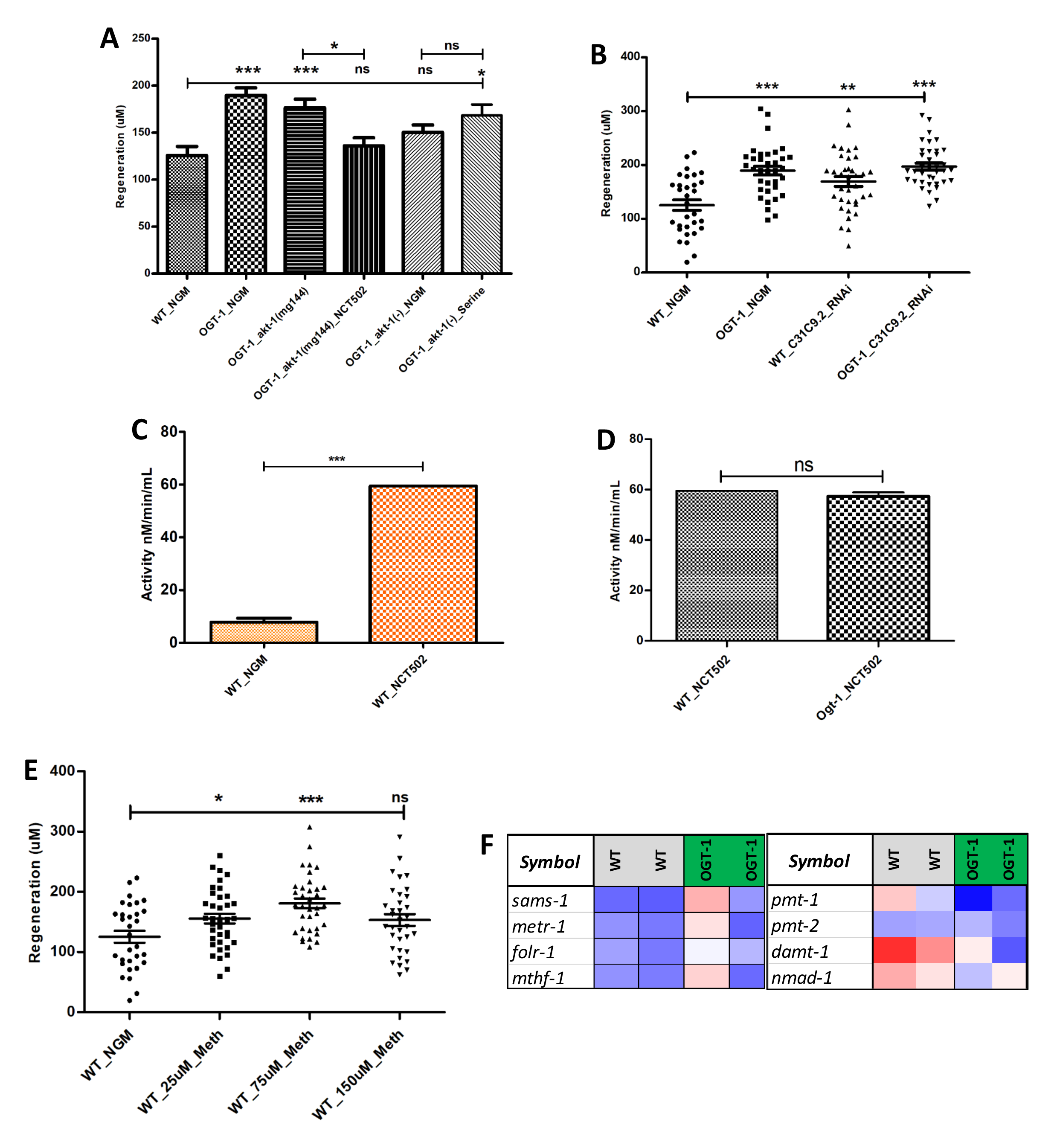
**(A)** The effects of NCT502 mediated inhibition of the serine synthesis pathway and serine supplementation on regeneration in *akt-1* (gain of function) and *akt-1* (loss of function) mutations in the *ogt-1* background. **(B)** 24 h neuron regeneration with systemic RNAi knockdown against C31C9.2 (ortholog of human PHGDH). **(C)** *pyk-1* activity in WT worms grown with and without NCT502 treatment. **(D)** *pyk-1* activity in WT and *ogt-1* worms grown with NCT502 treatment. **(E)** The effect of different doses of methionine supplementation on 24 h neuron regeneration in WT worms. **(F)** Expression, patterns of selected genes involved in One Carbon Metabolism (*sams-1, metr-1, folr-1 & mthf-1*), Transmethylation (*damt-1 & nmad-1*) and lipogenesis (*pmt-1 & pmt-2*) in neuronal cell RNAseq analysis which passed FDR 0.1. All data shown in ±SEM, analytical methods; student t-test and One Way ANOVA were used; ns, no significance; *pValue <0.05, **pValue <0.01, ***pValue <0.001.

**Supplemental Figure 5.**
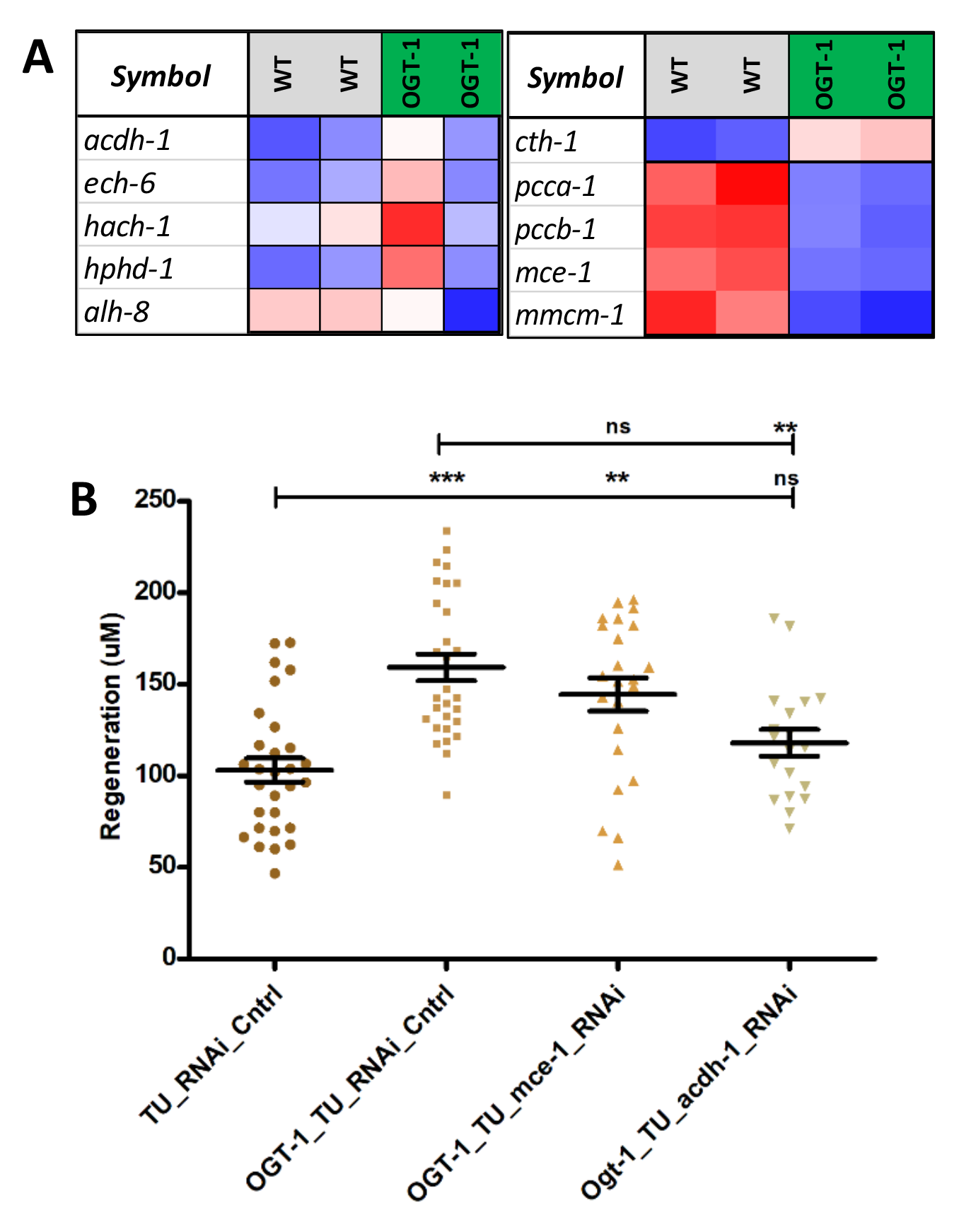
**(A)** Expression patterns, of selected genes involved in vitamin B12 independent shunt pathway (*acdh-1, each-6, hach-1, hphd-1 & alh-8*) and vitamin B12 dependent canonical pathway (*cth-1, pcca-1, pccb-1, mce-1 & mmc-1*) downstream to Transsulfuration Pathway (TSP), in neuronal cell RNAseq analysis which passed FDR 0.1. **(B)** The effect on 24 h neuron regeneration from neuron specific RNAi knock down of *acdh-1* and *mce-1* in *ogt-1* and WT worms. All data shown in ±SEM, analytical methods; One Way ANOVA was used; ns, no significance; *pValue <0.05, **pValue <0.01, ***pValue <0.001.

### Supplemental Tables

Table S1. Regeneration data for Fig. 1.

Table S2. Systemic RNAseq_DEGs Data.

Table S3. Neuronal RNAseq_DEGs Data.

Table S4. Regeneration data for Fig. 3.

Table S5. Regeneration data for Fig. 4.

Table S6. Regeneration data for supplemental Fig. 4 and Fig. 5.

Table S7. List of qRT-PCR primers used in the study.

